# STI1 domains coordinate partitioning of UBQLN2 into stress-induced condensates

**DOI:** 10.64898/2026.04.01.715099

**Authors:** William Haws, Thuy P. Dao, Bridget Varner, Holly B. Jones, Mallory P. Brown, Carlos A. Castañeda

## Abstract

UBQLN2 is a ubiquitin-binding shuttle protein that undergoes phase separation *in vitro* and localizes to stress-induced cellular condensates including stress granules. The central region of UBQLN2 contains two chaperone- and substrate-binding STI1 domains (STI1-I, STI1-II) and disordered linkers; the individual contributions of these domains and linkers to cellular condensate partitioning remain poorly characterized. Here we use live-cell imaging and immunofluorescence experiments to systematically examine domain requirements for UBQLN2 puncta formation in cultured human cells. We show that *in vitro* phase separation propensity largely correlates with puncta formation in transfected cells. Importantly, STI1-II and UBA domains are each required for baseline puncta formation in cells, but not STI1-I. In contrast, both STI1 domains are required for heat stress-induced puncta formation. Removal of STI1-II abrogates this stress response, and STI1-I deletion substantially attenuates it. Using N-terminal truncation constructs, we demonstrate that STI1-I strongly promotes both phase separation and puncta formation in the absence of the N-terminal region containing the UBL domain. Together, our findings demonstrate that the two STI1 domains of UBQLN2 have distinct roles in puncta formation and condensate partitioning, with STI1-II essential under all conditions.

**Highlights:**

- UBQLN2 is recruited to both stress granules and puncta formed by the autophagy receptor protein p62 in response to heat stress.
- Both endogenous and overexpressed UBQLN2 tend to colocalize with p62 puncta in cells where p62 is abundant, even without acute stress treatment.
- Using an extensive domain deletion library, ability of UBQLN2 constructs to form puncta correlates with their *in vitro* phase separation propensity in the absence of stress.
- Each of the two STI1 domains contribute non-redundantly to formation of UBQLN2 puncta in response to heat stress.
- The STI1-II domain is independently required for UBQLN2 oligomerization, phase separation, and puncta formation.

## Introduction

Ubiquilin-2 (UBQLN2) is a ubiquitin-binding shuttle protein that participates in multiple protein quality control (PQC) pathways, including substrate stabilization and degradation that invoke proteasomal degradation or autophagy (Hjerpe *et al*, 2016; Zheng *et al*, 2020; Zientara-Rytter & Subramani, 2019; Onwunma *et al*, 2026; Takei *et al*, 2025). The protein contains an N-terminal ubiquitin-like (UBL) domain that interacts with the proteasome and a C-terminal ubiquitin-associated (UBA) domain that engages ubiquitinated substrates (**Figure 1A**). While the functions of these folded domains are well-established, less is known about the middle region of UBQLN2 (residues 109-576). This central region contains two clusters of STI1 (stress-inducible protein 1) repeat motifs: STI1-I (residues 178-247) and STI1-II (residues 379-462) (Mah *et al*, 2000; Kleijnen *et al*, 2000). Uniquely among the UBQLN family in humans, this central region also harbors a proline-rich (PXX) domain. The majority of *UBQLN2* mutations linked to familial amyotrophic lateral sclerosis (ALS) and frontotemporal dementia (FTD) are in the PXX region, although several disease-linked mutations exist in the STI1 domains and the disordered linkers between the STI1 domains (Deng *et al*, 2011; Williams *et al*, 2012; Renaud *et al*, 2019; Lin *et al*, 2022). While only a small percentage of cases are directly caused by mutations in the corresponding gene, UBQLN2 is found within insoluble protein inclusions in diseased neurons of ALS patients regardless of etiology, frequently together with ubiquitinated substrates and the autophagy receptor p62 (Deng *et al*, 2011; Williams *et al*, 2012; Thumbadoo *et al*, 2024).

**Figure 1.**
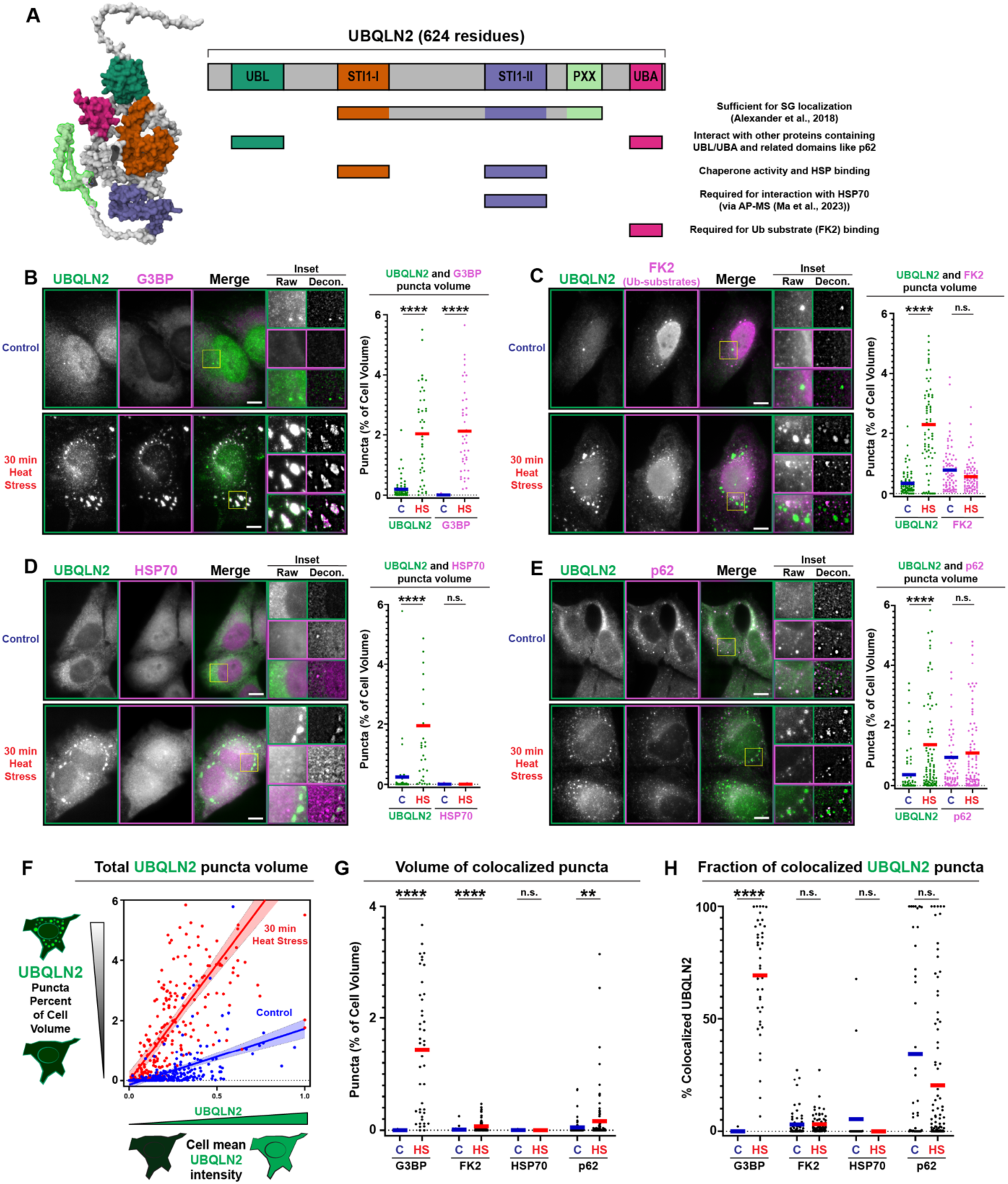
Endogenous UBQLN2 puncta formation correlates with protein quality control markers during heat stress. **(A)** Schematic of UBQLN2 domain architecture (AlphaFold: AF-Q9UHD9-F1-v6) and established motifs involved in interaction and/or colocalization with relevant PQC factors. Labeled domains are UBL (ubiquitin-like), STI1-I, STI1-II, PXX (proline-rich), and UBA (ubiquitin-associated). **(B–E)** Representative fluorescence images and puncta volume quantifications from HeLa cells subjected to 37°C control or 30 min 42°C heat stress, then immunolabeled for endogenous UBQLN2 (green) and G3BP **(B)**, FK2 (conjugated ubiquitin) **(C)**, HSP70 **(D)**, or p62 **(E)**. Scale bars = 10 μm. When paired with G3BP or FK2, rabbit anti-UBQLN2 (Sigma-Aldrich, HPA006431, 1:500) was used. When paired with p62 or HSP70, mouse anti-UBQLN2 (Novus NBP2-25164, 1:500) was used. Insets show raw and deconvolved signal for UBQLN2 (solid green border), respective co-labels (solid magenta border), and merged channels (gradient border). Graphs display the proportion of analyzed cell volume classified as puncta for UBQLN2 and the respective co-label. Horizontal bars represent the mean across all cells. C, control; HS, heat stress. **(F)** Overall UBQLN2 puncta percentage of cell volume plotted as a function of normalized mean UBQLN2 cell fluorescence intensity, with data pooled from experiments in **(B–E)**. Control (blue) and 30 min heat stress (red) conditions are shown. **(G)** The volume of UBQLN2 puncta that colocalize with G3BP, p62, FK2, or HSP70 (B-E) expressed as a percentage of each analyzed cell as in (B-F). **(H)** The voxel-wise fraction of UBQLN2 puncta that colocalize with G3BP, p62, FK2, or HSP70 **(B-E)**. For all statistical comparisons: *p < 0.05, **p < 0.01, ***p < 0.001, ****p < 0.0001, Mann-Whitney test. UBQLN2 was analyzed in cells under control/heat stress (C/HS) experiments, respectively with G3BP n=59/43, FK2 n=74/77, HSP70 n=40/32, p62 n=68/97.

Previously, we and others showed that UBQLN2 undergoes phase separation (PS) *in vitro*, forming liquid-like biomolecular condensates at physiological temperature and salt concentrations (Dao *et al*, 2018) and forming stress-induced cytoplasmic condensates under arsenite stress (Riley *et al*, 2021; Alexander *et al*, 2018; Dao *et al*, 2018). From inspection of a C-terminal construct of UBQLN2, we determined that the molecular origins of UBQLN2 phase separation stem from multivalent interactions distributed across the protein involving the STI1-II, PXX, and UBA domains (Dao *et al*, 2018, 2019). Of these, the STI1-II domain is key as it drives UBQLN2 oligomerization (e.g., dimerization) and phase separation (Dao *et al*, 2018).

Further increasing the complexity of understanding structural and functional relationships of UBQLN2, the central region of UBQLN2 (containing STI1-I and STI1-II domains) also interacts with the UBL and UBA domains (Zheng *et al*, 2021). Not only do these folded domains drive important heterotypic interactions with protein quality control components, the UBL and UBA domains also engage in intramolecular interactions with each other, creating a network of contacts that we hypothesize tunes phase separation propensity (Zheng *et al*, 2021). Recent structural and biophysical studies have revealed that the STI1 hydrophobic groove in UBQLNs engages both exogenous client proteins and internal ‘placeholder’ sequences within UBQLNs (Onwunma *et al*, 2026; Acharya *et al*, 2026). In yeast Dsk2, which contains a single STI1 domain, intramolecular STI1-helix interactions are a primary driver of self-association and phase separation (Acharya *et al*, 2026), but how each of the two STI1 domains in UBQLN2 contributes to condensate formation in mammalian cells is poorly characterized. The central region of UBQLN2 is also key in localizing UBQLN2 to stress granules, transient ribonucleoprotein condensates that assemble in response to diverse cellular stressors and are thought to protect mRNAs and stalled translation complexes until stress is relieved (Protter & Parker, 2016). While the central STI1-containing region of UBQLN2 is sufficient for stress granule localization (Alexander *et al*, 2018), this recruitment requires the cofactors RTL8 and PEG10 and these components are only beginning to be characterized (Mohan *et al*, 2025).

UBQLN2 also colocalizes with the autophagy receptor p62/SQSTM1 (hereafter p62) in both physiological and pathological contexts. p62 is itself a phase-separating protein that constitutively forms cytoplasmic droplets serving as hubs for selective autophagy (Kageyama *et al*, 2021; Sun *et al*, 2018; Zaffagnini *et al*, 2018). Unlike stress granules, which form transiently in response to acute stress, p62 puncta are present in unstressed cells and can evolve into larger, more gel-like “p62 bodies” under conditions of proteotoxic burden (Kageyama *et al*, 2021). UBQLN2 and p62 also colocalize into protein-containing inclusions in cellular and animal models of ALS (Deng *et al*, 2011; Thumbadoo *et al*, 2024), but p62 droplets and SGs appear to be mutually exclusive compartments (Chitiprolu *et al*, 2018; Turco *et al*, 2019). The molecular determinants that govern UBQLN2 recruitment to these distinct condensate populations remain undefined.

Here we systematically examine the relationship of UBQLN2 puncta formation in the context of protein quality control components, including p62. Furthermore, we determine the domain requirements for UBQLN2 puncta formation in cultured human cells without any experimental stress treatment (which we will refer to as “baseline” conditions) and in cells subjected to heat stress. Using a library of deletion constructs, we correlate results from both *in vitro* phase separation experiments on purified proteins and cell-based heat stress experiments with fluorescently-tagged UBQLN2 proteins. We uncover distinct contributions of the N-terminal region of UBQLN2 and of each STI1 domain to puncta formation and highlight differences in their respective contributions between baseline and heat stress conditions.

## Results

### Endogenous UBQLN2 puncta formation correlates with PQC markers during heat stress

To first establish the cellular contexts in which endogenous UBQLN2 forms puncta and to understand its relationships with other PQC proteins, we examined endogenous UBQLN2 in HeLa cells together with key interaction partners via paired immunofluorescence experiments (**Figure 1B-E**). We analyzed fixed and immunolabeled cells to measure the overall amount and colocalization of puncta formed by endogenous UBQLN2 and respective co-labeled proteins in control or heat stress conditions (cells heated to 42°C for 30 minutes). We focused on four potential colocalization protein partners: the core stress granule protein G3BP (**Figure 1B**), conjugated ubiquitin detected by FK2 (**Figure 1C**), HSP70-class heat shock proteins (**Figure 1D**), and the autophagy receptor p62 (**Figure 1E**). We quantified UBQLN2 puncta volume separately in each paired comparison and consistently identified a small population of UBQLN2 puncta in unstressed cells. Thirty minutes of heat stress significantly increased the volume of UBQLN2 puncta relative to control across all four pairings (**Figure 1B-E**). Cells with higher mean UBQLN2 antibody fluorescence intensity, used here as a proxy for relative protein concentration, were more punctate than lower-intensity cells in both control and heat stress conditions (**Figure 1F**). To assess colocalization between UBQLN2 puncta and each co-label, we identified voxels where respective puncta overlapped and quantified them both as a proportion of cell volume (**Figure 1G**) and as a fraction of total UBQLN2 puncta per cell (**Figure 1H**).

Consistent with past studies, stress granules labeled with G3BP were virtually absent from unstressed cells but were robustly induced by 30 min of heat stress (**Figure 1B**). It is well-established (Dao *et al*, 2018; Alexander *et al*, 2018; Mohan *et al*, 2025; Phung *et al*, 2023) that heat stress induces formation of stress granules that recruit UBQLN2. As expected, we observed a substantial amount of UBQLN2-positive stress granules (**Figure 1G**), and these constituted the majority of overall UBQLN2 puncta in this condition (**Figure 1H**).

Conjugated ubiquitin detected by FK2 also localized to discrete puncta in both control and heat stress conditions, with a similar volume of FK2-labeled puncta between treatments (**Figure 1C**). However, FK2 puncta were largely mutually exclusive of UBQLN2 puncta in either condition. While the volume of overlapping puncta increased slightly with heat stress (**Figure 1G**), this was proportional to the increase in overall UBQLN2 puncta with no change in fractional colocalization (**Figure 1H**).

In contrast to the other analyzed proteins, HSP70 rarely localized to discrete puncta in either control or heat stress conditions (**Figure 1D**), and the small amount of HSP70 puncta had little overlap with those of UBQLN2 (**Figure 1G, 1H**). HSP70 was not actively excluded from UBQLN2-labeled puncta consistent with stress granules (**Figure 1D inset**). HSP70 did not concentrate in stress granules or other cytoplasmic structures in response to this short heat stress treatment.

We next examined UBQLN2 together with the autophagy receptor p62 (**Figure 1E**). This protein constitutively localized to puncta in control conditions, and the per-cell puncta proportion was not significantly altered by 30 min of heat stress (**Figure 1E**). Uniquely among this set of co-labeled proteins, a significant fraction of UBQLN2 puncta in control cells colocalized with p62 (**Figure 1H**). The volume of colocalized puncta was increased by heat stress (**Figure 1G**) proportional to the overall increase in UBQLN2 puncta, but with similar fractional colocalization between the two conditions (**Figure 1H**). On average, both the volume and fraction of colocalized UBQLN2 puncta were lower for p62 than for G3BP under heat stress conditions (**Figure 1G, 1H**). Whereas UBQLN2 colocalization with stress granules in heat-stressed cells was relatively consistent, the amount of colocalized UBQLN2/p62 puncta varied substantially between cells in both control and heat stress conditions (**Figure 1H**).

### Overexpressed UBQLN2 is recruited to p62 droplets

To investigate the factors driving recruitment of UBQLN2 to cellular stress-induced puncta, we generated an mEos3.2-tagged UBQLN2 fusion construct for transient transfection. We subjected cells overexpressing mEos3.2-UBQLN2 to identical stress and fixation treatments as in our endogenous immunolabeling experiments (**Figure 1**). In this case, we used retained mEos3.2 fluorescence to detect exclusively the overexpressed protein in one channel and immunofluorescence for the co-labeled endogenous protein in the other (**Figure S1A, S1B**). Like endogenous UBQLN2, mEos3.2-UBQLN2 formed puncta under control conditions, and the amount of puncta was increased with 30 min of heat stress (**Figure 2A, 2B**). However, paired immunofluorescence detection of G3BP revealed only mild recruitment of mEos3.2-UBQLN2 to stress granules (**Figure 2A, 2C, 2D**), consistent with our and others’ prior reports of poor recruitment of overexpressed UBQLNs to stress granules (Alexander *et al*, 2018; Dao *et al*, 2018; Riley *et al*, 2021).

**Figure 2.**
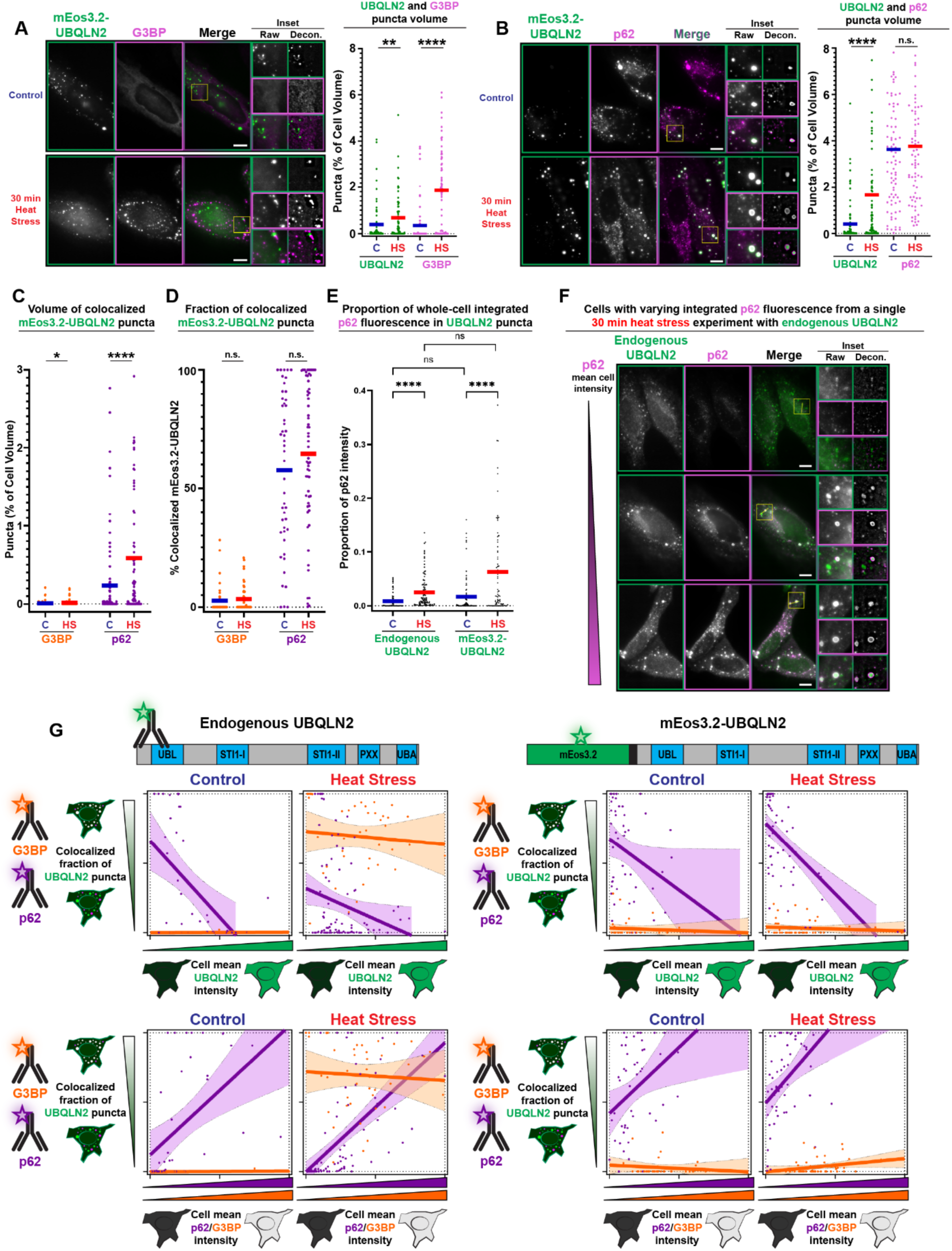
UBQLN2 is recruited to p62 puncta. **(A,B)** Representative images and puncta volume quantifications from HeLa cells transfected with mEos3.2-UBQLN2 (green), subjected to 37°C control or 30 min 42°C heat stress, then immunolabeled for endogenous G3BP **(A)** or p62 **(B)**. Scale bars = 10 μm. Insets show raw and deconvolved signal. Graphs display the proportion of analyzed cell volume classified as puncta for mEos3.2-UBQLN2 and the respective co-label individually, as well as the voxel-wise proportion of analyzed cell volume where puncta overlap. Horizontal bars represent the mean across all cells. C, control; HS, heat stress. **(A-D)** *p < 0.05, **p < 0.01, ***p < 0.001, ****p < 0.0001, Mann-Whitney test. **(C)** The volume of mEos3.2-UBQLN2 puncta that colocalize with G3BP or p62. **(D)** The voxel-wise fraction of mEos3.2-UBQLN2 puncta that colocalize with G3BP or p62. **(E)** The proportion of whole-cell integrated p62 fluorescence intensity contained within spatially delimited UBQLN2 puncta, quantified per cell for endogenous UBQLN2 and mEos3.2-UBQLN2 experiments under control and heat stress conditions. Horizontal bars represent the mean across all cells. C, control; HS, heat stress. ns, not significant; ****p < 0.0001, Kruskal–Wallis test with Dunn’s multiple comparisons correction. **(F)** Three representative HeLa cells from a single 30 min heat stress experiment immunolabeled for endogenous UBQLN2 (green) and p62 (magenta), shown in ascending order of mean p62 cell intensity. Scale bars = 10 μm. Insets show raw and deconvolved signal. **(G)** Colocalized fraction of UBQLN2 puncta plotted as a function of normalized cell mean intensities of respective targets for endogenous UBQLN2 (left) and mEos3.2-UBQLN2 (right), shown under control and heat stress conditions. Top row: colocalized fraction as a function of normalized UBQLN2 cell mean intensity. Bottom row: colocalized fraction as a function of normalized partner (p62 or G3BP) cell mean intensity. Data points and regression lines with shaded 95% confidence intervals are shown for p62 (purple) and G3BP (orange). mEos3.2-UBQLN2 was analyzed under control/heat shock (C/HS) experiments with G3BP n=65/71 and p62 n=83/74 cells.

We next assessed colocalization with p62 (**Figure 2B**). As with endogenous UBQLN2, mEos3.2-UBQLN2 puncta had a large degree of overlap with those of p62 even in control conditions (**Figure 2D**). Similarly, the volumes of total mEos3.2-UBQLN2 puncta and colocalized puncta both increased with heat stress (**Figure 2B, 2C**) without a concomitant increase in overall p62 puncta volume (**Figure 2B**). We reasoned that stress induced additional recruitment of UBQLN2 to a relatively invariant population of p62 puncta. For both endogenous and overexpressed UBQLN2 protein, we found that heat stress increased the fraction of total cellular p62 fluorescence intensity contained within voxels designated as UBQLN2 puncta (**Figure 2E**), consistent with recruitment of UBQLN2 to a volume of p62 that remained constant between control and heat stress conditions. In summary, mEos3.2-UBQLN2 was poorly recruited to stress granules, but shared similar patterns of p62 colocalization as endogenous UBQLN2 protein (compare **Figure 2B** and **2F**).

Using mean cellular fluorescence intensity as a proxy for protein abundance, we also found that cells with higher amount of UBQLN2 contained more UBQLN2 puncta (**Figure 1F**). We next asked whether similar correlations existed between overall protein abundance and puncta colocalization between respective pairs of fluorescently-labeled target proteins for both endogenous and overexpressed UBQLN2 (**Figure 2G**). We treated the fraction of colocalized UBQLN2 puncta in each cell as a function of the mean cell fluorescence intensity of immunolabeled endogenous UBQLN2 or G3BP and found no significant relationship (**Figure 2G** - left). This was also true for mEos3.2-UBQLN2 paired with G3BP, even in low-expressing cells (**Figure 2G** - right). However, colocalization with p62 was highly correlated with abundance for both endogenous and overexpressed UBQLN2 in both control and heat stress conditions. Interestingly, in each case colocalization of puncta with p62 was negatively correlated with UBQLN2 intensity but positively correlated with p62 intensity (**Figure 2G**). This was not due to an increase in overall UBQLN2 puncta in high-p62 cells, as we found no such association between mean cell p62 intensity and total UBQLN2 puncta volume (**Figure S2**). This contrasts with G3BP, FK2, and HSP70, the mean cell fluorescence intensities of which positively predicted UBQLN2 puncta formation under heat stress in paired experiments (**Figure S2**).

### UBQLN2 domains differentially contribute to puncta formation in living cells

While heat-induced recruitment to stress granules was low for mEos3.2-UBQLN2 relative to the endogenous protein, they shared similar patterns of p62 recruitment and expression-dependent puncta formation in both control and heat stress conditions (**Figure 2**). We therefore used the mEos3.2-UBQLN2 construct and targeted deletions thereof in live-cell imaging experiments to systematically identify domains of UBQLN2 involved in puncta formation. We first focused on systematically removing each domain individually, including the folded domains UBL (1-108) and UBA (577-620), the two STI1 domains STI1-I (178-247) and STI1-II (379-462), and the disease-linked PXX (491-538) region **(Figure 3A)**. We transfected HEK293 cells with these mEos3.2-fusion constructs and imaged them 24-30 hours post-transfection with no added stress treatment (baseline) (**Figure 3**). We acquired images at a single, focus-locked Z-plane, segmented cells and puncta in 2D, and calculated the respective fraction of pixels classified as puncta for each cell (**Figure 3B, Figure S1C**). To facilitate comparisons across constructs and minimize the potential for overexpression artifacts, we analyzed cells below an arbitrarily chosen mean cell intensity threshold (see Methods). Full-length mEos3.2-UBQLN2 formed puncta under baseline conditions that totaled about 0.5% of segmented cell area on average (**Figure 3B**). Puncta formation was variable across the different domain deletion constructs ranging from 0 to <2% of cell area. Puncta formation was expression-dependent as we identified a statistically significant relationship between puncta area and the mean fluorescence intensity for mEos3.2-UBQLN2 and deletion constructs thereof (**Figure 3D**), but not mEos3.2-only. These results are fully consistent with previously published reports on exogenous expression of UBQLN2 (Deng *et al*, 2011; Sharkey *et al*, 2018; Dao *et al*, 2018).

**Figure 3.**
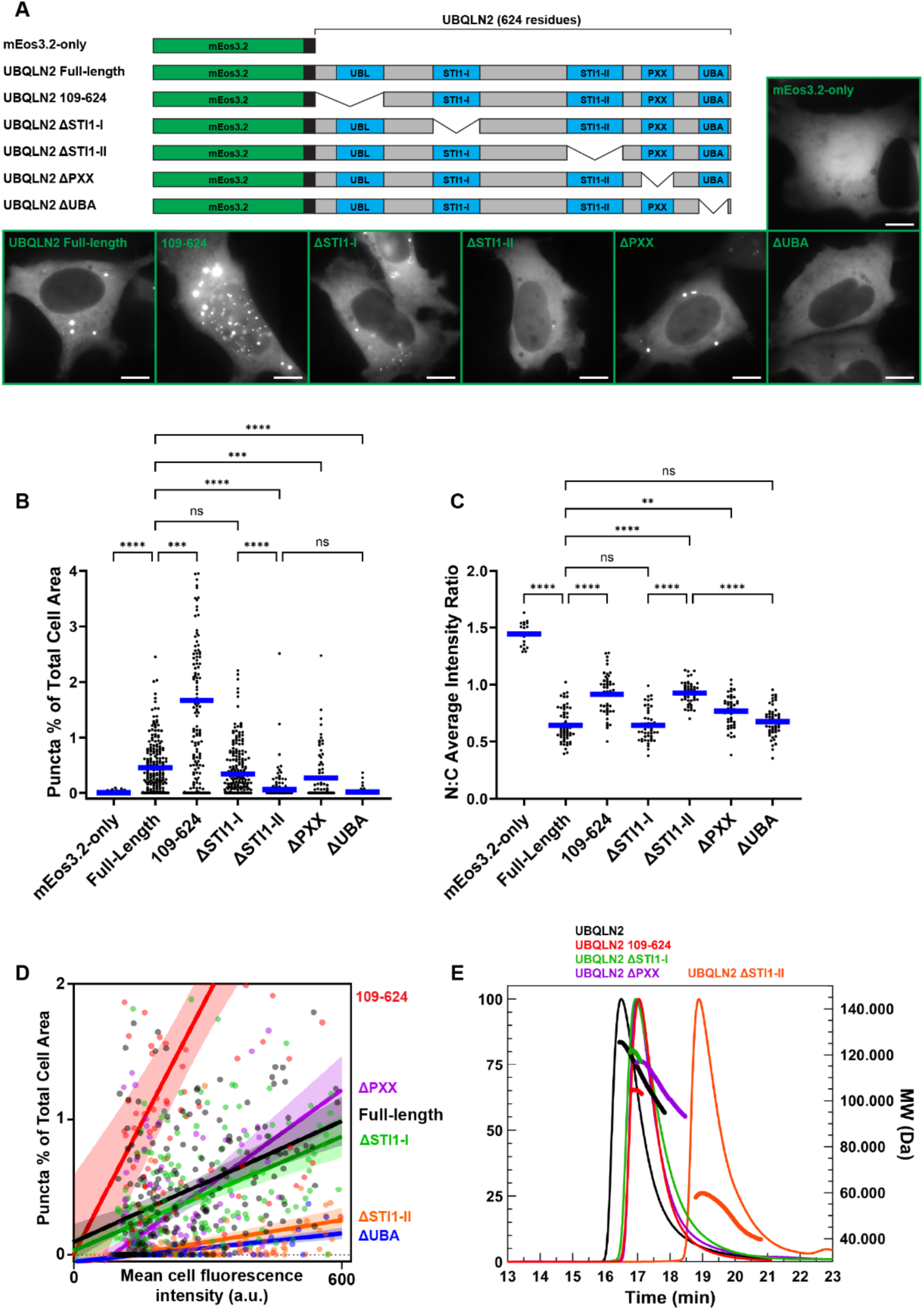
UBQLN2 puncta formation is modulated by its constituent domains. **(A)** Schematic representations of mEos3.2-UBQLN2 constructs and representative images of HEK293 cells imaged 24–30 hours post-transfection. Scale bars = 10 μm. **(B)** Quantification of the percent area of each cell classified as puncta from a single focal plane. Data points reflect individual cells (n >= 62 per construct) from pooled experiments (n=6). Blue bars represent mean values across all cells in all experiments. *p<0.05, **p<0.01, ***p<0.001, ****p<0.0001 Kruskal-Wallis Test with Dunn’s Multiple Comparisons Correction. **(C)** Nuclear:cytoplasmic (N:C) average fluorescence intensity ratios derived from manually traced nuclear boundary annotations in a subset of cells (n = 19 for mEos3.2-only, n >= 48 per UBQLN2 construct) from pooled experiments (n = 6). Blue bars represent mean values. ns, not significant; **p < 0.01, ****p < 0.0001, Kruskal–Wallis test with Dunn’s multiple comparisons correction. **(D)** Puncta area percentage (same data presented in (B)) plotted as a function of the mean fluorescence intensity of the corresponding cell. Linear regression curves and confidence intervals were calculated per-construct and are shown color-coded along with individual data points. **(E)** Size-exclusion chromatography with multi-angle light scattering (SEC-MALS) analysis of the indicated UBQLN2 constructs expressed and purified from *E. coli* without fusion partners or epitope tags.

### N-terminal UBL deletion increases puncta formation

Phase separation of purified UBQLN2 *in vitro* is strongly enhanced by removal of the first 108 residues (Zheng *et al*, 2021), which constitute the ∼30-residue N-terminal IDR and the subsequent UBL domain. Cellular puncta formation of a mEos3.2-UBQLN2 construct lacking these residues (109-624) was significantly enhanced relative to the full-length protein (**Figure 3B**), consistent with previous results that show an increased puncta formation of UBQLN2 109-624 in neurons (Sharkey *et al*, 2018). We further showed that expression levels did not explain the difference in puncta formation vs. the full-length protein for this or any other deletion construct, as we found identical relationships after normalizing to mean cell fluorescence intensity (**Figure S3**). We also noted that UBQLN2 109-624 robustly formed both nuclear and cytoplasmic puncta, unlike full-length. We delineated nucleus vs. cytoplasm (N:C) puncta in a subset of cells via manual tracing of the apparent N:C boundary (**Figure S4**). UBQLN2 109-624 formed significantly more puncta in the nucleus than full-length UBQLN2. From quantification of the gross fluorescence ratio inside vs. outside nuclei, UBQLN2 109-624 had a significantly higher N:C ratio than full-length UBQLN2 (**Figure 3C**).

### Deletion of either STI1-II or UBA domain abrogates most baseline puncta formation

The central region of UBQLN2 contains two STI1 domains that contribute both to self-interactions and to interactions with chaperones and client proteins (Hjerpe *et al*, 2016; Kurlawala *et al*, 2017; Onwunma *et al*, 2026; Itakura *et al*, 2016). While deletion of the first STI1 domain (ΔSTI1-I) had no discernable effect relative to full-length UBQLN2, deletion of STI1-II abrogated almost all puncta formation in transfected cells (**Figure 3B**). This loss of puncta was not due to competition with endogenous UBQLN2, as mEos3.2-UBQLN2-ΔSTI1-II similarly formed almost no puncta in UBQLN2 knockout cells (**Figure S5**). The ΔSTI1-II construct also had an elevated N:C fluorescence intensity ratio relative to both full-length and ΔSTI1-I (**Figure 3C**). While the overall increase in N:C ratio was very similar to that of UBQLN2 109-624, nuclear ΔSTI1-II fluorescence was diffuse. Removal of the UBA domain (ΔUBA) likewise nearly eliminated puncta formation under baseline conditions (**Figure 3B**). While ΔSTI1-II and ΔUBA were both diffusely distributed, ΔUBA was more cytoplasmic and did not have an altered N:C ratio compared to full-length UBQLN2 (**Figure 3C**).

UBQLN2 also contains a PXX domain that distinguishes it from other UBQLN paralogs in humans and is the site of many ALS-causative mutations. Deletion of PXX reduced puncta formation in baseline conditions (**Figure 3B**) and increased N:C ratio (**Figure 3C**) relative to the full-length construct, though in each case these changes were mild compared to deletion of STI1-II.

We hypothesized that the molecular origins of baseline UBQLN2 puncta formation in cells were related to UBQLN2’s propensity for oligomerization. We previously determined that UBQLN2 dimerization was a driving factor in providing the multivalency needed for UBQLN2 phase separation under physiological conditions *in vitro* (Dao *et al*, 2018, 2019). Therefore, we analyzed the oligomerization propensity of purified domain deletion constructs using size-exclusion chromatography coupled to multi-angle light scattering (SEC-MALS) (**Figure 3E**). Strikingly, all tested domain deletion constructs were dimeric except for ΔSTI1-II, which was monomeric. We conclude that the STI1-II domain is independently required both for oligomerization *in vitro* and for puncta formation in baseline conditions, while the STI1-I domain is dispensable for both phenomena. The exception to the above is ΔUBA, which we and others previously determined to be dimeric using native mass spectrometry (Sharkey *et al*, 2018; Robb *et al*, 2023). We also showed that ΔUBA has a reduced propensity to phase separate *in vitro* (Zheng *et al*, 2021), consistent with what we observed in cells.

### STI1 domains drive formation of UBQLN2 puncta in response to heat stress

Like endogenous UBQLN2, we identified a greater volume of full-length mEos3.2-UBQLN2 puncta in cells fixed after a 30 min heat stress treatment than in control conditions (**Figure 2A**, **2B**). We next used live-cell, time lapse experiments to quantify heat stress-induced puncta formation (**Figure 4, Movie S1**). HEK293 cells expressing mEos3.2-UBQLN2 constructs were imaged once every 10 minutes for two hours (a total of 13 timepoints). Initial timepoint images were taken at 37°C, after which the temperature was shifted to 42°C for heat stress or kept at 37°C (control). Full-length mEos3.2-UBQLN2 puncta consistently comprised about 0.5% of segmented cell area through all timepoints of control experiments (**Figure 4A**, 4B). About 30 minutes after increasing the temperature, mEos3.2-UBQLN2 formed additional puncta that plateaued at an average of 1.2% of cell area (**Figure 4A, 4B**).

**Figure 4.**
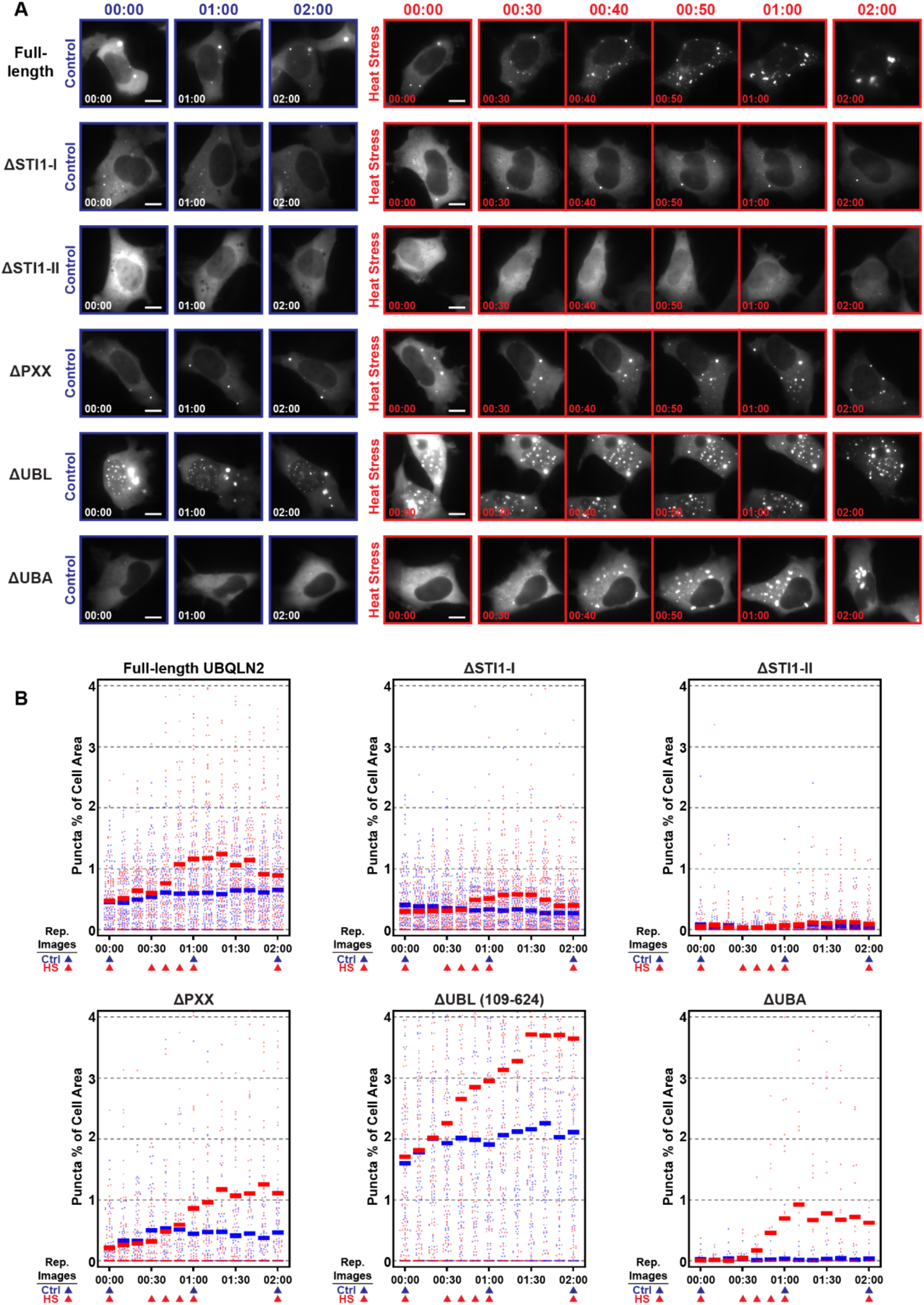
STI1 domains drive formation of UBQLN2 puncta in response to heat stress. **(A)** Representative live-imaging time series of HEK293 cells transfected with full-length mEos3.2-UBQLN2, ΔSTI1-I, ΔSTI1-II, ΔPXX, 109-624 (lacks the UBL domain), or ΔUBA. Cells were subjected to either control (37°C) or heat stress (ramp from 37°C to 42°C) conditions. Images were acquired every 10 min for 2 hours. Control cells are shown at 0, 1, and 2 hours. Heat stress cells are shown at 0, 30, 40, 50 min, and 1 and 2 hours, capturing the window during which most *de novo* puncta formation occurred. Scale bars = 10 μm. **(B)** Quantification of the percent of total cell area classified as puncta at all timepoints for each construct. Data points represent individual cells from control (blue) and heat stress (red) experiments (n = 3 each). Horizontal bars represent mean values across all cells. The number of analyzed cells at outset were as follows for respective control and heat stress (C/HS) timelapse experiments: Full-Length: n=98/124; ΔSTI1-I: n=116/141; ΔSTI1-II: n=111/105; ΔPXX: n=102/66; 109-624: n=96/101; ΔUBA: n=29/24.

To identify the domains of UBQLN2 specifically required for heat stress-induced puncta formation, we subjected cells expressing various deletion constructs to this timelapse paradigm (**Figure 4A, Figure S6**). The ΔPXX constructs formed heat stress-induced puncta at a more gradual level than full-length UBQLN2. The 109-624 construct lacking the UBL formed the largest percentage of heat stress-induced puncta (nearly 4% of cell area). By contrast, the ΔUBA construct (which was almost completely diffuse in initial timepoints and remained so through the duration of control experiments) readily formed stress-induced puncta in cells subjected to heat stress (**Figure 4A, 4B**). The timing and amount of *de novo* puncta formation were both similar to that of full-length UBQLN2. In contrast, this response was attenuated for each of the independent STI1 deletions (ΔSTI1-I or ΔSTI1-II in **Figure 4A, 4B**). While similar to full-length at baseline, ΔSTI1-I only reached about 0.6% puncta at any point during heat stress (**Figure 4B**). Removal of STI1-II had an even stronger effect; ΔSTI1-II showed almost no puncta formation in response to heat stress, increasing only to about 0.1% at later stages (**Figure 4B**).

In these live-cell experiments, ΔSTI1-II was the only mEos3.2-UBQLN2 construct we assessed that showed no change in puncta formation either in control or heat stress conditions over time (**Figure 4B**). We also fixed and immunolabeled endogenous p62 in ΔSTI1-II-transfected cells and similarly found no change in ΔSTI1-II puncta between control and 30-minute heat stress treatments (**Figure S7A, S7B**), consistent with the lack of heat stress-induced puncta formation in living cells. Unlike endogenous UBQLN2 (**Figure 1G**) or full-length mEos3.2-UBQLN2 (**Figure 2C**), heat stress did not increase the volume of p62-colocalized puncta (**Figure S7B**) or lead to enrichment of ΔSTI1-II in individual p62 puncta (**Figure S7C**). Together, these data suggest non-redundant roles for the respective STI1 domains in driving heat-induced formation of UBQLN2 puncta, compared to control conditions.

### *In vitro* UBQLN2 phase separation trends correlate with UBQLN2 puncta formation in cells

In addition to the stress-specific roles of the STI1 domains, our single domain deletion analysis revealed a correlation between cellular puncta formation and *in vitro* phase separation experiments. Enhancement or inhibition of cellular puncta formation in the absence of stress by respective deletion of UBL or UBA domains corresponded closely to changes in phase separation that we characterized previously (Zheng *et al*, 2021; Dao *et al*, 2018). We aimed to systematically compare phase separation propensity with cellular UBQLN2 puncta formation.

We created a set of successive N-terminal deletion constructs, assessed their *in vitro* phase separation propensity, cellular puncta formation in baseline conditions, and acute puncta formation in response to heat stress. We quantified the phase separation propensity of different UBQLN2 constructs in terms of cloud point temperature and saturation concentration (c_sat_). For all constructs, phase separation was observed with increasing temperature, consistent with LCST (lower critical solution temperature) behavior as previously observed for UBQLN2. We obtained temperature-concentration phase diagram data for this set of N-terminal deletions and several domain deletion constructs (**Figure 5A**). Notably, ΔSTI1-II did not phase separate in the concentration and temperature ranges tested. To compare ΔSTI1-II directly with full-length, we performed sedimentation assays in the presence of PEG8000 as a crowding agent (**Figure S8**). These conditions enhanced phase separation of full-length UBQLN2, reducing its c_sat_ to only ∼6 µM at 35°C. Under these conditions, the concentration threshold for ΔSTI1-II phase separation remained very high at ∼100 µM.

**Figure 5.**
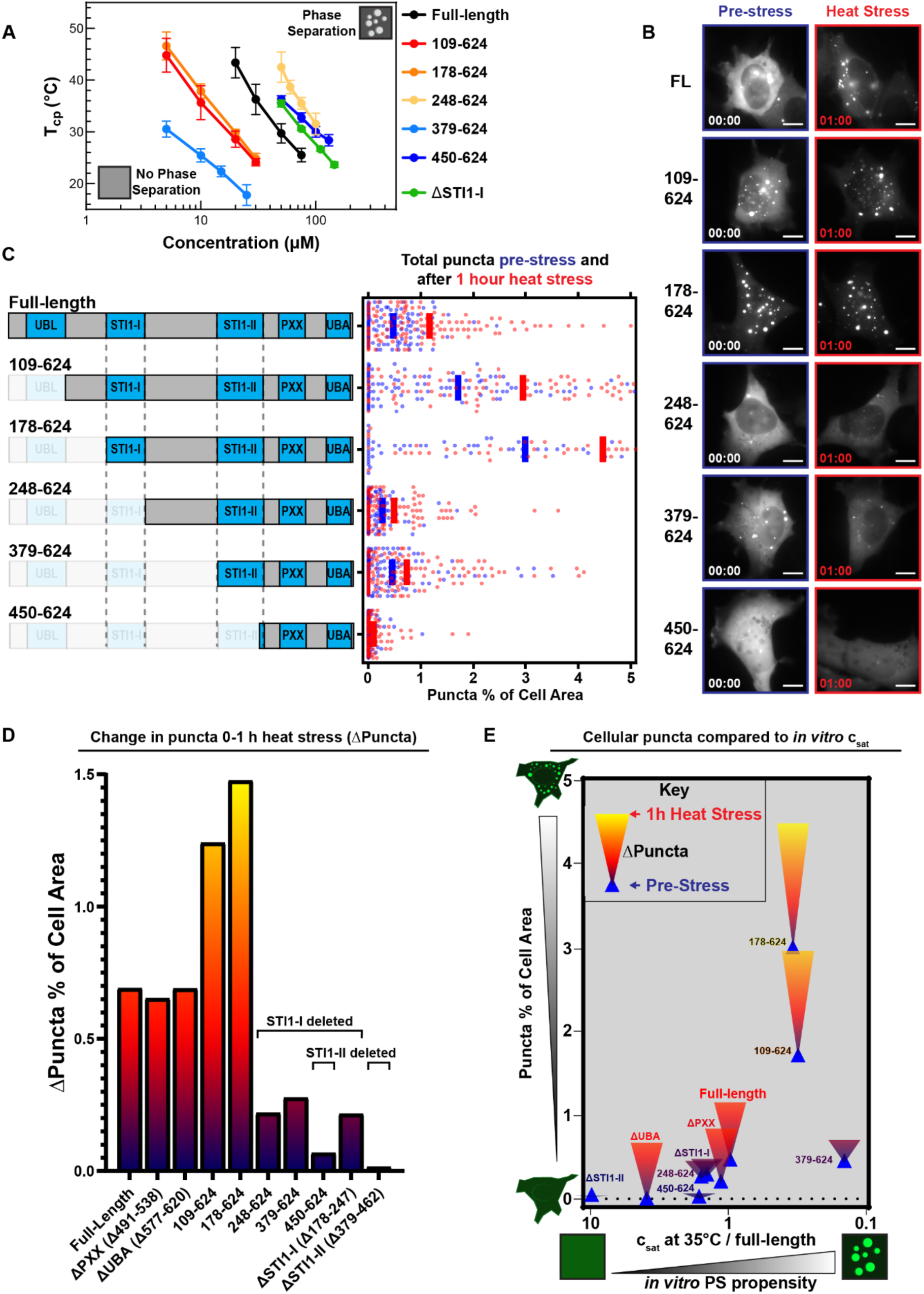
The STI1-I domain has important roles in UBQLN2 localization and stress response that are constrained by STI1-UBL interactions. **(A)** Cloud-point temperatures (T_cp_) as a function of protein concentration for the indicated UBQLN2 constructs, determined by turbidity assays with proteins expressed and purified from *E. coli* without fusion partners or epitope tags in pH 6.8 buffer containing 200 mM NaCl. ΔSTI1-II is not shown as it did not phase separate under these conditions. Data points represent mean ± SEM across replicates. **(B-C)** Representative images **(B)** and per-cell puncta area fractions **(C)** pre-stress (blue data points and bar representing mean) and after 1 hour 42°C heat stress (red data points and bar representing mean) for each construct. Full results for all constructs are shown in Figure S6. **(D)** Change in mean puncta area between pre-stress and 1 hour heat stress timepoints (ΔPuncta) for each construct. **(E)** The change in mean puncta area between pre-stress and 1 h heat stress timepoints (ΔPuncta) for each construct plotted against saturation concentrations (c_sat_) derived from *in vitro* phase separation assays (**(A)**, ΔPXX from (Dao *et al*, 2024), and ΔUBA from (Zheng *et al*, 2021)) with matching constructs. The c_sat_ values plotted here are divided by the c_sat_ value of full-length UBQLN2 at 35°C. The ΔSTI1-II c_sat_ value is positioned at the bottom left edge of the plot given its very low phase separation propensity (Figure S8). Blue triangles = pre-stress puncta percentage of cell area; inverted triangles = ΔPuncta spanning pre-stress and 1 hour heat stress averages and colored according to maximum bar height in **(D)**. For the comparison of relative saturation concentration against pre-stress puncta: Spearman rho = −0.84, p = 0.002.

Sequential removal of N-terminal segments from UBQLN2 had variable effects on the promotion and inhibition of phase separation compared to full-length UBQLN2. Deletion of the first 177 residues that include the UBL and disordered linker (UBQLN2 109-624, 178-624 constructs) promoted phase separation of UBQLN2 as the phase diagrams shifted to the left. However, further removal of the STI1-I region (UBQLN2 248-624) dramatically inhibited phase separation of UBQLN2. Strikingly, expression of these constructs in cells followed the same pattern: both 109-624 and 178-624 readily formed puncta in both the nucleus and cytoplasm under baseline conditions (**Figure 5A, 5B, S6**), but UBQLN2 248-624 had reduced propensity to form puncta compared to full-length UBQLN2 (**Figure 5B**). Interestingly, both UBQLN2 248-624 (lacking the UBL and STI1-I) and the independent removal of the STI1-I region (ΔSTI1-I) had similar effects *in vitro* and in cells (**Figure 3B, 5B**). Collectively, these results suggest that the N-terminal region of UBQLN2 interacts with the STI1-I domain and/or that the 1-248 region of UBQLN2 has variable interactions with other components in the cell to regulate its promotion or inhibition of UBQLN2 puncta formation.

As heat-induced UBQLN2 puncta formation was attenuated by deletions of either STI1 domain (**Figure 4**), we tested whether we would observe the same patterns in the N-terminal truncation constructs by using our live-imaging heat stress paradigm (**Figure 4, S6**). We quantified the change in puncta (as a proportion of cell area) from pre-stress to 1 hour of heat stress for each construct (**Figure 5B, 5C**) and compared this heat stress-induced change (ΔPuncta) (**Figure 5D**) to pre-stress puncta and *in vitro* phase separation propensity (**Figure 5E**). We found that those constructs with increased phase separation propensity were the best at forming cellular puncta under either baseline conditions or heat stress (**Figure 5E**).

Remarkably, those UBQLN2 constructs that contained *both* STI1-I and STI1-II domains produced the largest change in heat-induced puncta formation (first five in **Figure 5D**). Heat stress response was attenuated for those constructs lacking the STI1-I (e.g., UBQLN2 248-624, 379-624, ΔSTI1-I). Those constructs that lacked most of or the entire STI1-II domain (e.g., UBQLN2 450-624, ΔSTI1-II) were the least responsive in forming heat stress-induced puncta. Taken together, these data lead us to conclude that the STI1 domains autonomously and non-redundantly drive formation of UBQLN2 puncta in response to acute heat stress (**Figure 6**).

**Figure 6.**
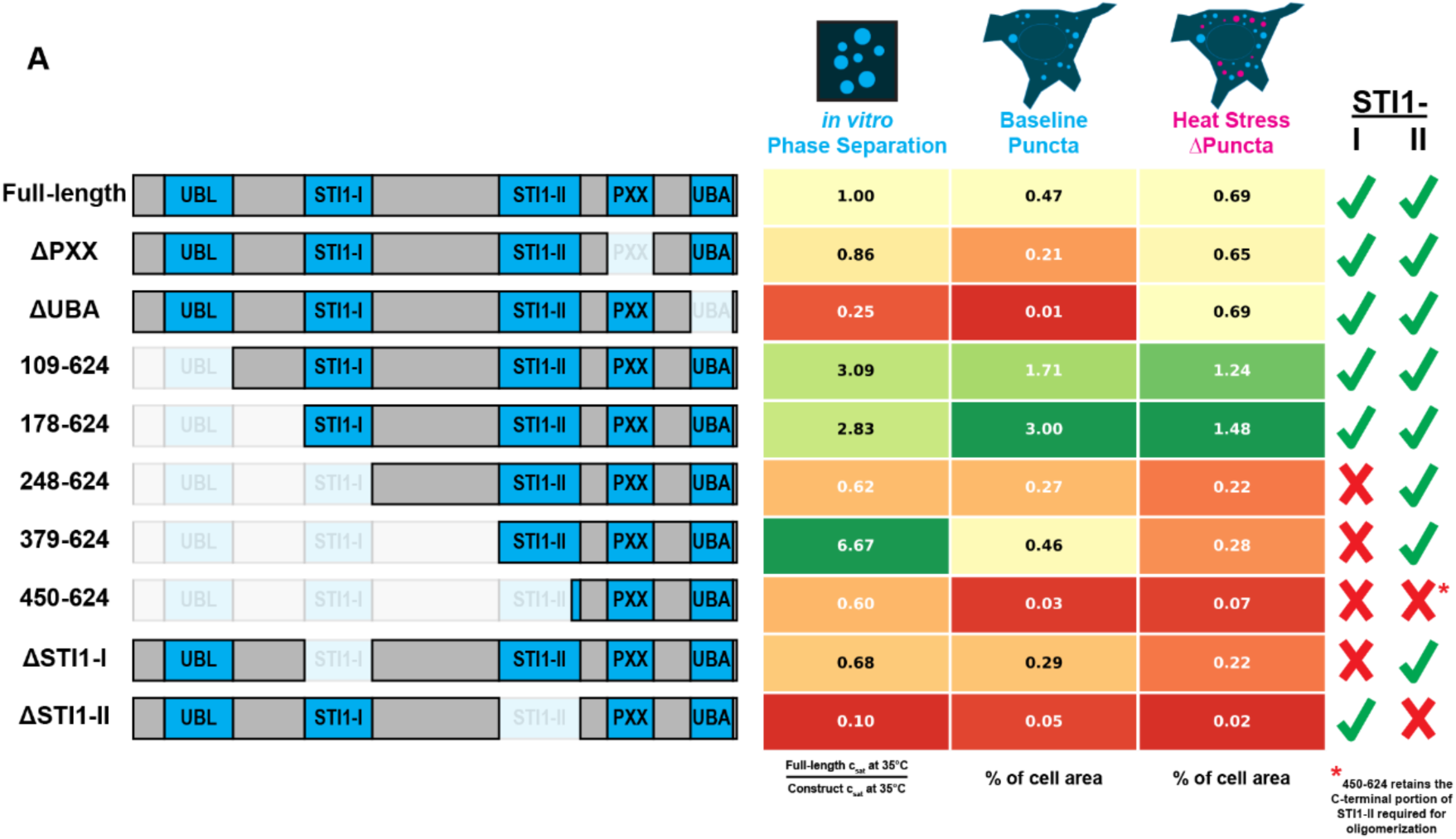
Summary table of UBQLN2 construct properties. Domain schematics are shown alongside normalized *in vitro* phase separation propensity (full-length c_sat_ at 35°C / construct c_sat_ at 35°C), baseline puncta (% of cell area in cells without added stress), heat stress ΔPuncta (change in % of cell area). Values for full-length UBQLN2 are shaded as yellow boxes, with increased (high) and decreased (low) values relative to full-length UBQLN2 colored green and red, respectively. Right-hand side indicates whether the designated construct contains (green checkmark) or lacks (red X) the STI1-I and STI1-II domains.

## Discussion

In this study, we demonstrated that endogenous UBQLN2 localized to heat-induced stress granules as previously shown, but more importantly, we observed correlations between UBQLN2 puncta volume and the cellular levels of several protein quality control markers that did not themselves localize to the same puncta. Heat-induced UBQLN2 puncta formation and stress granule localization correlated with increased HSP70 and FK2 (marking ubiquitinated substrates) fluorescence intensities (**Figure 1, S2**), suggesting that UBQLN2 puncta formation also responds to proteostatic burden rather than co-aggregating with these factors. This pattern suggests that elevated HSP70 and ubiquitination levels may promote UBQLN2 puncta formation through transient interactions rather than stable co-recruitment to the same puncta. This interpretation is consistent with the hypothesis that UBQLN2 acts as a sensor of ubiquitinated substrate load. As we show that UBQLN2 puncta formation in cells correlates with UBQLN2 phase separation trends *in vitro* (**Figure 5**), we speculate that UBQLN2 cellular puncta may also be responding to changes in ubiquitin/polyubiquitin levels (Dao *et al*, 2018, 2022; Galagedera *et al*, 2023; Valentino *et al*, 2024).

While overexpressed UBQLN2 showed poor recruitment to stress granules, consistent with prior reports (Alexander *et al*, 2018; Riley *et al*, 2021), it robustly associated with p62 puncta under both control and heat stress conditions (**Figure 2A**). The proportion of cellular p62 contained within UBQLN2 puncta increased approximately three-fold with heat stress, while the overall volume and number of p62 puncta remained relatively constant (**Figure 2E**). This pattern suggests that heat stress promotes UBQLN2 recruitment to a pre-existing population of p62 puncta rather than driving formation of new p62 structures. Indeed, endogenous UBQLN2 also is recruited to p62 puncta, particularly at higher expression levels of p62 (**Figure 2F**). Interestingly, p62 colocalization fraction was positively correlated with p62 intensity but negatively correlated with UBQLN2 intensity (**Figure 2G**). One explanation for this asymmetry is that with higher UBQLN2 expression, a greater proportion of the protein partitions into homotypic UBQLN2 condensates driven by STI1-II-mediated self-association, effectively competing with heterotypic co-condensation at p62 puncta. The co-condensation of UBQLN2 and p62 is significant due to their shared presence in pathological inclusions in ALS patients carrying *UBQLN2* mutations (Deng *et al*, 2011). Unlike stress granule markers such as G3BP, which are not found in disease-associated aggregates, p62 is a consistent component of UBQLN2-positive inclusions in both familial and sporadic ALS (Thumbadoo *et al*, 2024; Wu *et al*, 2020; Deng *et al*, 2011; Williams *et al*, 2012). Our observation that these proteins co-condense in living cells under stress conditions suggests a potential route through which physiological stress responses could, if dysregulated, give rise to pathological deposits.

*In vitro* UBQLN2 phase separation propensity, as measured by saturation concentration (c_sat_ values), correlated strongly with baseline puncta formation across constructs (Spearman rho = −0.84, p = 0.002) (**Figure 5**). Constructs with enhanced phase separation propensity, such as those lacking the N-terminal UBL domain (109-624, 178-624), formed substantially more puncta, while constructs with severely impaired phase separation (ΔSTI1-II, ΔUBA) were almost completely diffuse under baseline conditions (**Figure 3B, 5A, S6**).

The cell response to heat stress revealed a more complex picture that highlighted the distinct functional roles of the two STI1 domains in UBQLN2. Deletion of STI1-II virtually eliminated both baseline and stress-induced puncta formation, consistent with its essential role in driving UBQLN2 self-association and phase separation. In contrast, deletion of STI1-I had no detectable effect on baseline puncta but substantially attenuated the heat stress response (**Figure 4B**). This asymmetry suggests that STI1-I makes contributions to UBQLN2 condensation that are latent in the full-length protein under baseline conditions and become functionally relevant during heat stress. We speculate that these contributions occur through mechanisms that are distinct from, and potentially additive to, the homotypic interactions mediated by the STI1-II domain (**Figure 6**).

The most striking demonstration of STI1-I function came from our comparative analysis of successive N-terminal deletions. UBQLN2 178-624, which retains STI1-I but lacks the UBL domain and preceding sequences, displayed high phase separation propensity and formed abundant cellular puncta. Deletion of STI1-I (resulting in UBQLN2 248-624) dramatically reduced both *in vitro* phase separation and puncta formation in cells (**Figure 5A-D**), suggesting that STI1-I makes substantial positive contributions to phase separation. However, since the isolated STI1-I deletion from full-length UBQLN2 had minimal effects (compare ΔSTI1-I with full-length in **Figure 3**), these positive contributions are masked in the full-length protein by N-terminal UBL-mediated autoinhibition. This interpretation is in agreement with our prior evidence from NMR spectroscopy for direct interactions between 1-108 (containing the UBL) and the middle part of UBQLN2 including STI1-I (Zheng *et al*, 2021). Furthermore, we recently demonstrated that the single STI1 domain of yeast UBQLN Dsk2 engages transient amphipathic helices within its intrinsically disordered regions to drive self-association and phase separation (Acharya *et al*, 2026). The STI1-I domain of UBQLN2 shares similarity with the Dsk2 STI1 domain, raising the possibility that analogous intramolecular STI1-helix interactions contribute to UBQLN2 condensation when the N-terminal autoinhibitory region is removed or displaced. We speculate that when the 1-108 region is removed, STI1-I becomes available to participate in additional homotypic or heterotypic interactions that enhance phase separation. Notably, UBQLN2 109-624 and 178-624 constructs also displayed elevated nuclear localization and formed puncta in both nuclear and cytoplasmic compartments, suggesting that the N-terminus may additionally influence subcellular distribution, potentially through interacting with STI1-I domains. An intriguing implication is that cellular factors that bind the UBL domain, such as proteasomal subunits, could in principle relieve autoinhibition and promote UBQLN2 condensation. This mechanism would provide a way to couple UBQLN2 phase behavior to proteasomal engagement, potentially coordinating substrate sequestration with degradation pathway status.

The STI1-II domain was essential for baseline puncta formation in cells and for heat stress. The ΔSTI1-II construct formed almost no puncta in living cells under either baseline or heat stress conditions (**Figure 3B, 4A, 4B**). It has also been reported that STI1-II, but not STI1-I, is required for UBQLN2 to interact with HSP70 proteins (Ma *et al*, 2023); notably, this was an affinity purification experiment where both bait and prey proteins were tagged and overexpressed. Deletion of STI1-II also perturbed colocalization with p62 droplets as assessed by paired immunofluorescence (**Figure S7B, S7C**). Because ΔSTI1-II is nearly completely diffuse, the absence of colocalization with p62 is expected and does not by itself demonstrate a specific role for STI1-II in p62 recruitment; rather, it indicates that UBQLN2 condensation (which requires STI1-II) is a prerequisite for partitioning into the p62 droplet phase. In the few cells that did contain clear ΔSTI1-II puncta, their distribution and morphology were also notably different from those formed by full-length and other constructs (**Figure S6A**), though we did not systematically investigate this phenomenon. The STI1-II domain may contribute to puncta formation through multiple mechanisms: by driving UBQLN2 oligomerization and increasing its effective valency for co-phase separation with other components such as p62, or by enabling interactions with client substrates that promote condensation (Valentino *et al*, 2024; Acharya *et al*, 2026).

The STI1 domains of UBQLNs have recently been recognized as sites of client protein engagement. The STI1 hydrophobic groove binds to internal “placeholder” sequences within UBQLN2, including the PXX region, and ALS mutations disrupt these interactions (Onwunma *et al*, 2026). This model suggests that STI1-mediated intramolecular contacts contribute to the multivalency that drives phase separation, and that client binding can competitively displace placeholder sequences to regulate condensate behavior. Our observation that PXX deletion mildly reduced baseline puncta formation (**Figure 3B**) is consistent with the PXX region contributing to STI1-mediated self-association. Additionally, it was recently shown that UBQLN2 condensates catalyze alpha-synuclein fibril formation with alpha-synuclein binding to the STI1-II region (Takei *et al*, 2025). Finally, while most ALS-linked UBQLN2 mutations cluster in the PXX region, several map to the STI1 domains and linker regions (Deng *et al*, 2011; Williams *et al*, 2012; Renaud *et al*, 2019; Lin *et al*, 2022). Our finding that STI1-II is essential for all puncta formation while STI1-I contributes to stress-responsive puncta provides a framework for understanding how mutations in these domains could differentially affect basal versus stress-induced UBQLN2 condensation. Together with our finding that STI1-II is essential for all forms of UBQLN2 condensation, these studies position the STI1 domains as nodes linking UBQLN2 phase behavior, client engagement, and pathological aggregation.

## Conclusions

Together, our data establish that the two STI1 domains of UBQLN2 coordinate condensate partitioning through distinct, non-redundant mechanisms. STI1-II provides the oligomerization capacity essential for all forms of UBQLN2 condensation, while STI1-I makes latent contributions that become functionally relevant under heat stress when the autoinhibitory influence of the N-terminal region is partially overcome. These findings extend recent structural and functional studies that STI1 domains are client-engagement sites in UBQLNs (Acharya *et al*, 2026; Onwunma *et al*, 2026; Takei *et al*, 2025). UBQLN2 condensation and client engagement are likely mechanistically coupled through the STI1 domains. How client binding, chaperone interactions, and additional modulators such as post-translational modifications interact to regulate this coupling in physiological and disease contexts remains an important question for future investigation.

## Materials and Methods

### Cell Culture

HEK293 cells (ATCC CRL-1573) and HeLa cells (ATCC CCL-2) were maintained in Dulbecco’s Modified Eagle Medium (DMEM) supplemented with 10% fetal bovine serum (FBS) and 1% penicillin/streptomycin at 37 °C and 5% CO_2_. Cells were used at passage numbers below 35. UBQLN2 knockout HEK293 cells were a kind gift from the Lisa Sharkey and Hank Paulson laboratories at the University of Michigan. Cell lines were not independently authenticated.

### Plasmid Construction

The mEos3.2-UBQLN2 mammalian expression construct was derived from the mCherry-UBQLN2 plasmid described previously (Dao *et al*, 2018; Riley *et al*, 2021). Domain deletion and N-terminal truncation constructs were generated by Gibson assembly. The following domain deletion constructs were produced, with residue numbering referring to full-length human UBQLN2 (UniProt: Q9UHD9, 624 residues): ΔSTI1-I (lacking residues 178-247), ΔSTI1-II (lacking residues 379-462), ΔPXX (lacking residues 491-538), and ΔUBA (lacking residues 577-620, retaining residues 621-624). The ΔUBL construct was generated by deletion of residues 1-108, yielding a protein beginning at residue 109 (construct 109-624). N-terminal truncation constructs 178-624, 248-624, 379-624, and 450-624 were generated in the same backbone. The mEos3.2-only control construct consisted of the mEos3.2 coding sequence and linker region without any UBQLN2 sequence. All constructs were verified by whole-plasmid nanopore sequencing (Plasmidsaurus). For bacterial expression, UBQLN2 constructs were cloned into expression vectors by Gibson assembly as described previously (Dao *et al*, 2024).

### Transient Transfection Experiments

Cells were seeded onto glass-bottom imaging dishes (MatTek P35G-1.5-14-C; 35 mm dish, No. 1.5 coverslip, 14 mm glass diameter, uncoated) and transfected with 1 µg (HEK) or 2 µg (HeLa) DNA using TransIT-LT1 Transfection Reagent (MirusBio MIR 2300) according to the manufacturer’s protocol. Cells were imaged or fixed 24-30 h post-transfection.

### Heat Stress Treatment

For live-cell timelapse experiments, temperature was controlled using the built-in temperature-regulated stage of the ONI Nanoimager (Oxford Nanoimaging Ltd.). The stage temperature was ramped from 37 °C to 42 °C over the course of approximately 1 h, and cells were maintained at 42 °C through the remainder of the 2 h imaging period. Control cells were maintained at 37 °C throughout. For fixed-cell immunofluorescence experiments, heat stress was applied by transferring dishes to an incubator pre-warmed to 42 °C for 30 min, after which cells were immediately fixed.

### Live-Cell Fluorescence Microscopy

HEK cells were imaged in Leibovitz’s L-15 Medium, no phenol red (ThermoFisher 21083027). All imaging was performed on an ONI Nanoimager equipped with a Hamamatsu sCMOS ORCA Flash 4.0 V3 camera and an Olympus 100x/1.4 NA oil-immersion objective. The pixel size was 117 nm x 117 nm. The field of view was approximately 50 x 80 µm per acquisition. For live-cell imaging of mEos3.2-tagged constructs, the unconverted (green) form of mEos3.2 was excited using the 473 nm laser at 14% power, and emission was collected through the lower-wavelength channel of the instrument’s dichroic mirror. A single focus-locked Z-plane was acquired for each field. Ten frames of 100 ms exposure each were captured per field and averaged to produce the final image used for analysis. A total of 120 fields were acquired per live-cell imaging grid. All imaging parameters were held constant across experiments and constructs to enable direct quantitative comparison. For timelapse heat stress experiments, fields were revisited once every 10 min for 2 h (13 total timepoints). The first timepoint was acquired at 37 °C before initiation of the temperature ramp. A visual description of our imaging protocol is provided in **Figure S1**.

### Immunofluorescence

Cells were simultaneously permeabilized and partially fixed for 10 min at room temperature in a solution of 0.1% Triton X-100 and 2% paraformaldehyde (PFA) in phosphate-buffered saline (PBS), followed by a 20 min fixation step in 2% PFA in PBS. Fixed cells were blocked for 1 h at room temperature in 5% donkey serum in PBS supplemented with 0.25% Tween-20 (PBST). Primary antibodies were diluted in PBST + 3% w/v BSA and incubated overnight at 4 °C. The following primary antibodies were used: Mouse anti-UBQLN2 (Novus NBP2-25164, 1:500), Rabbit anti-UBQLN2 (Sigma-Aldrich, HPA006431, 1:500), Mouse anti-G3BP1/2 (BD Biosciences 611127, 1:500), Rabbit anti-p62/SQSTM1 (Proteintech 18420-1-AP, 1:500), Rabbit anti-HSP70 (Proteintech 10995-1-AP, 1:500), FK2 Mouse anti-conjugated ubiquitin (Enzo, LSI-AB-0120, 1:500). After three washes in PBS, cells were incubated with fluorophore-conjugated secondary antibodies (Donkey anti-Rabbit Alexa Fluor 647 (Invitrogen A-31573, 1:500), Donkey anti-Rabbit Alexa Fluor 488 (Invitrogen A-21206, 1:500), Donkey anti-Mouse Alexa Fluor 647 (Invitrogen A-31571, 1:500) as appropriate) diluted in PBST + 3% w/v BSA for 1 hour at room temperature. After three additional PBS washes, cells were stored in fresh PBS in the original MatTek dishes at 4°C prior to imaging. For colocalization experiments with overexpressed mEos3.2-UBQLN2, retained native mEos3.2 fluorescence in the green channel was combined with immunofluorescence detection of endogenous proteins in the far-red channel.

### Fixed-Cell Imaging

Fixed immunofluorescence samples were imaged on the same ONI Nanoimager described above. Z-stacks of 9 slices spaced 400 nm apart were acquired, with each slice collected at 100 ms exposure. The 473 nm laser was used at 14% power for the mEos3.2 or green-channel signal, and the 640 nm laser was used at 7-8% power (depending on target) for the far-red immunofluorescence channel. Each fixed-cell image was analyzed as a single field without stitching. Where indicated for representative image insets, iterative deconvolution was performed using the Iterative Deconvolve 3D plugin in Fiji (ImageJ) (Schindelin *et al*, 2012, 2015). Deconvolution was applied only to displayed images and not to images used for quantitative analysis.

### Machine Learning-Based Image Segmentation

Trainable Weka Segmentation (Arganda-Carreras *et al*, 2017) classifiers were trained for automated segmentation of cells and puncta. Training was performed on representative images acquired under the same conditions as experimental data. The feature set included features from the ImageScience library in addition to the default Weka feature set. A single pair of classifiers (one for cell segmentation, one for puncta segmentation) was trained and applied uniformly to all mEos3.2-UBQLN2 live-cell imaging data without modification. For 3D immunofluorescence data, Weka 3D segmentation was applied to Z-stacks to segment puncta in each channel independently. Probability maps generated by the classifiers were thresholded using empirically determined, target-specific and model-specific threshold values that were held constant for a given target and imaging condition. Segmented puncta were filtered to include only objects of at least 9 pixels (each pixel 117 nm x 117 nm, corresponding to a minimum puncta area of approximately 0.12 µm^2^).

### Puncta Quantification and Analysis

For live-cell 2D analysis, the puncta area fraction was calculated for each segmented cell as (puncta pixel area) / (total cell pixel area). Mean fluorescence intensity was measured within each segmented cell. Only cells with mean mEos3.2 fluorescence intensity less than 600 arbitrary units (AU) above the camera read floor were included in analysis.

Nuclear-to-cytoplasmic (N:C) fluorescence ratios were calculated from a subset of cells in which the nuclear boundary was manually traced based on apparent contrast in the mEos3.2 fluorescence signal. The N:C ratio was computed as the ratio of mean fluorescence intensity inside versus outside the traced nuclear boundary.

### Colocalization Analysis

For 3D colocalization analysis of immunofluorescence data, puncta were independently segmented in each fluorescence channel as described above. Colocalization was assessed by identifying voxels where segmented puncta from the two channels overlapped spatially. The colocalized volume was quantified both as a proportion of the total analyzed cell volume and as a fraction of total UBQLN2 puncta volume per cell. Mean fluorescence intensity values were normalized to a 0-1 range using min-max normalization applied within pooled replicates of the same target pair.

### Recombinant Protein Expression and Purification

UBQLN2 constructs for *in vitro* biophysical characterization were expressed in *Escherichia coli* BL21(DE3) cells and purified without the use of affinity tags, following the UBQLN-specific purification protocol described previously (Dao *et al*, 2018, 2024). Briefly, bacterial cultures were grown in Luria-Bertani (LB) medium supplemented with appropriate antibiotics at 37 °C to an OD_600_ of approximately 0.6-0.8, induced with isopropyl beta-D-1-thiogalactopyranoside (IPTG), and incubated at reduced temperature to allow protein expression. Cell pellets were lysed and proteins were purified by a combination of ammonium sulfate precipitation, anion exchange chromatography, and size-exclusion chromatography (SEC), as described (Dao *et al*, 2018, 2024). Purified proteins were buffer-exchanged into 20 mM sodium phosphate, 200 mM NaCl, 0.5 mM TCEP, 1 mM EDTA (pH 6.8) unless otherwise noted. Protein purity was assessed by SDS-PAGE, and concentrations were determined by UV absorbance at 280 nm using calculated extinction coefficients.

### Size-Exclusion Chromatography Coupled with Multi-Angle Light Scattering (SEC-MALS)

The oligomeric state of purified UBQLN2 constructs was determined by SEC-MALS. These measurements were performed at BioCAT (beamline 18ID at the Advanced Photon Source, Chicago) using in-line size exclusion chromatography (SEC) to separate sample from aggregates and other contaminants thus ensuring optimal sample quality and multiangle light scattering (MALS), dynamic light scattering (DLS) and refractive index measurement (RI) for additional biophysical characterization (SEC-MALS-SAXS). The samples were loaded on a Superdex 200 Increase 10/300 Increase column (Cytiva) run by a 1260 Infinity II HPLC (Agilent Technologies) at 0.6 ml/min. The eluent passed sequentially through an Agilent UV detector, MALS detector (18-angle DAWN Helios II, Wyatt), SAXS sample cell, and RI detector (Optilab T-rEX, Wyatt). Molecular weights were calculated from MALS/RI data using ASTRA 7 (Wyatt).

### Phase Separation Assays

Temperature-concentration phase diagrams for all constructs, except for ΔSTI1-II, were constructed by measuring saturation concentrations (c_sat_) as a function of temperature. Phase separation of UBQLN2 constructs was assessed by turbidity and sedimentation assays as described previously (Dao *et al*, 2018, 2024; Galagedera *et al*, 2023). For turbidity assays, purified protein at indicated concentrations in 20 mM sodium phosphate, 200 mM NaCl, 0.5 mM TCEP, 1 mM EDTA (pH 6.8) was placed in a temperature-controlled UV-Vis spectrophotometer (Agilent 3500). Optical density at 600 nm (OD_600_) was monitored as a function of temperature during a controlled ramp. Cloud point temperatures were determined as the temperature at which OD_600_ exceeded a defined threshold above baseline.

We quantified the saturation concentration (c_sat_) values for the full-length and ΔSTI1-II constructs using SDS-PAGE gels (Figure S8). 15 μL of 2× stocks of UBQLN constructs (30 μM full-length, and 300 μM for ΔSTI1-II) were mixed with 15 μL of 2× salt solution (20 mM sodium phosphate (pH 6.8), 0.5 mM TCEP, 0.5 mM EDTA, 300 mM NaCl, 5% PEG8000) and incubated for 10 min in temperature-equilibrated centrifuges at 10°C, 20°C, 30°C, 40°C, and 50°C. Samples were then centrifuged for 5 min at 21,000 × *g*. 1 μL of the supernatant of each sample was immediately mixed with 20 μL of 2× SDS-PAGE dye. 4 μL of each sample was loaded onto 10% polyacrylamide gels, imaged using the BioRad Gel Doc EZ Imager, and band volumes were determined with BioRad Image Lab software. For each gel, samples for a standard curve containing the solution concentration for a particular experiment (c_sol_), c_sol_/2, c_sol_/4, c_sol_/8, c_sol_/16 μM of protein were also loaded, analyzed, and used to calculate the c_sat_ values of the proteins. The measurements were carried out using at least two different protein stocks with two trials for each.

### Statistical Analysis

All statistical analyses were performed using GraphPad Prism (version 10.6). Nonparametric tests were used throughout unless otherwise indicated. For comparisons of puncta area fractions and colocalization metrics between two conditions (e.g., control vs. heat stress), Mann-Whitney U tests were applied. For comparisons across multiple constructs, the Kruskal-Wallis test was used followed by Dunn’s post-hoc test for pairwise comparisons. For relationships between mean cell fluorescence intensity and puncta area fraction (e.g., Figure 3D), linear regression was performed using least-mean-squares fitting, and the significance of the regression was assessed by F-test. All statistical tests were two-sided unless otherwise noted. Sample sizes (number of cells and number of independent biological replicates) and exact p values are reported in figure legends.

## Acknowledgements

We acknowledge support from National Institutes of Health (NIH) R01 GM136946 and R35 GM158070 to CAC. We thank Dr. Maxwell Watkins for the SEC-MALS experiments performed at the BioCAT facility. This research also used resources of the Advanced Photon Source, a U.S. DOE Office of Science User Facility operated for the DOE Office of Science by Argonne National Laboratory under Contract No. DE-AC02-06CH11357. BioCAT was supported by grant P30 GM138395 from the National Institute of General Medical Sciences (NIGMS) of the NIH. We thank Dr. Anitha Rajendran, Dr. Nirbhik Acharya for several scientific discussions about this work. We acknowledge support from Alana Ramirez on data analysis, Gianna Frank for assistance on some cell-based experiments, and Sini Shqair for assistance on *in vitro* phase separation experiments. The content is solely the responsibility of the authors and does not necessarily reflect the official views of NIGMS or the NIH.

## Data and Code Availability

All raw imaging data, processed quantification data, and analysis code is deposited here: TBD

