## Supplementary Information for "STI1 domains coordinate partitioning of UBQLN2 into stress-induced condensates"

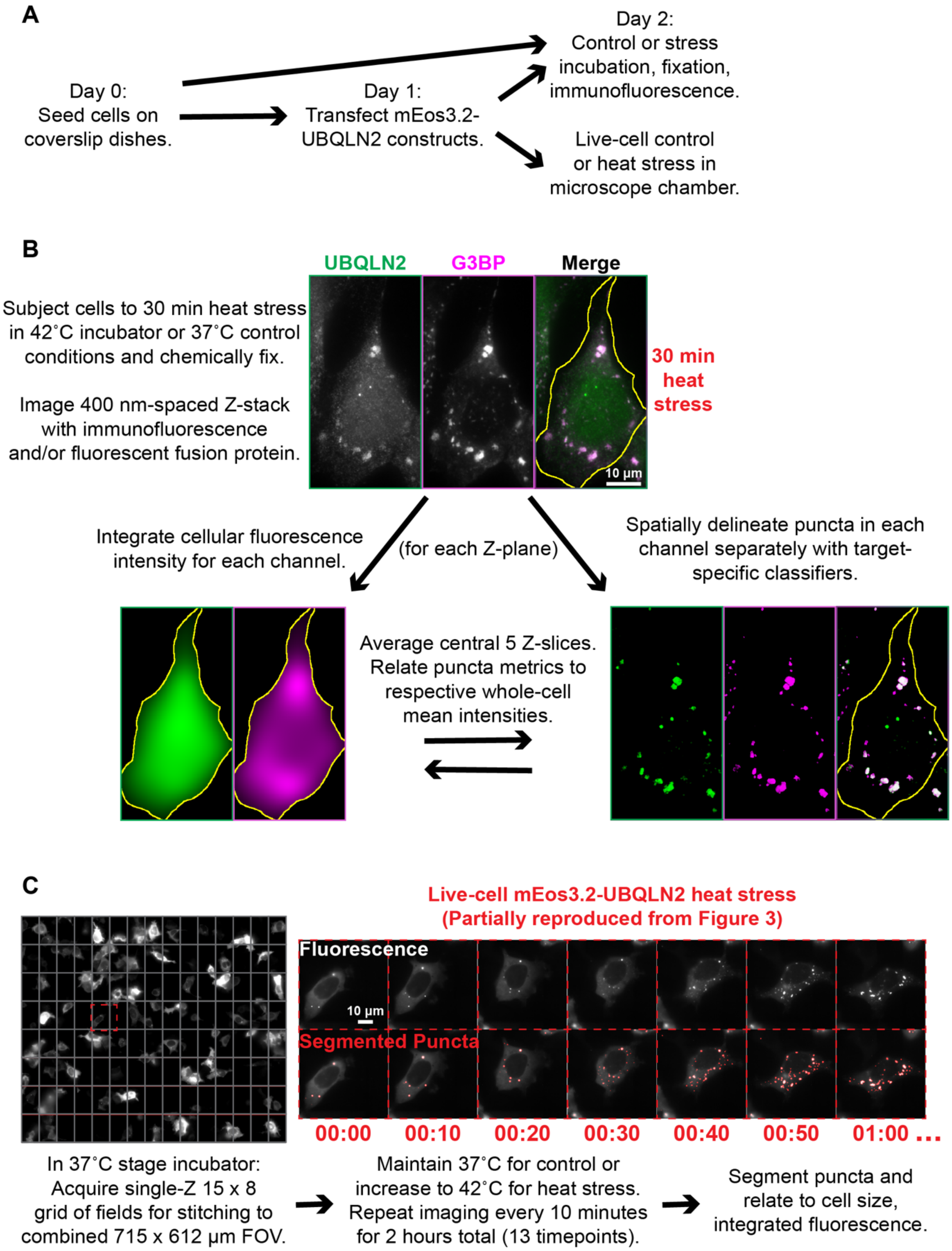


**Figure S1. Cell imaging and analysis workflow.**
(A) Experimental timeline for cell-based experiments. (B) Our protocol for handling fixed cell experiments is shown. (C) The protocol for live-cell experiments is shown (see **Movie S1**).

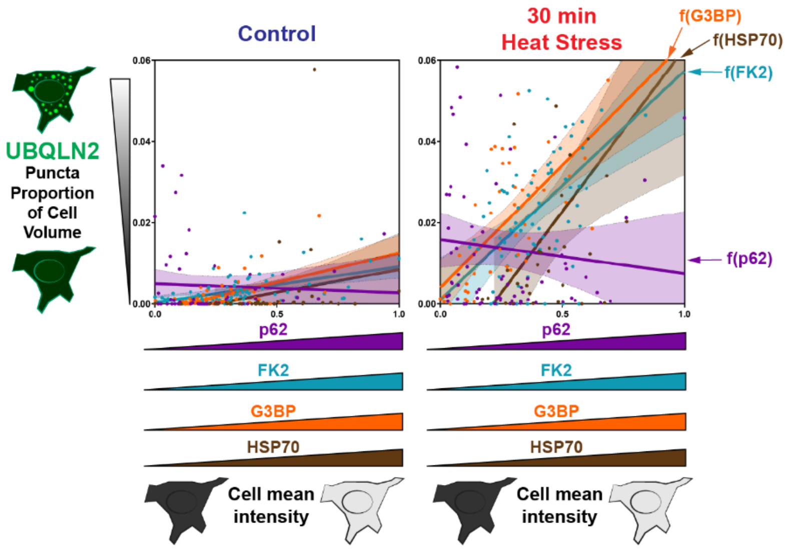


**Figure S2.** Endogenous UBQLN2 puncta formation as a function of co-label mean fluorescence cell intensity from experiments described in Figure 1. Mean fluorescence intensities normalized 0 to 1 within the combination of control and heat stress datasets for each co-label. Linear regression with 95% CI plotted. Analyzed number of cells from paired control/heat stress (C/HS) experiments with G3BP n=59/43, FK2 n=74/77, HSP70 n=40/32, p62 n=68/97 (C/HS). Spearman rho for G3BP = 0.3988 (C), 0.6638 (HS); FK2 = 0.5034 (C), 0.4362 (HS); HSP70 = 0.5013 (C), 0.5062 (HS); p62 = 0.1818 (C), -0.153 (HS). Spearman p values for control and heat stress with G3BP, FK2, HSP70 <0.01; for p62 control p = 0.1379, heat stress p = 0.1345.

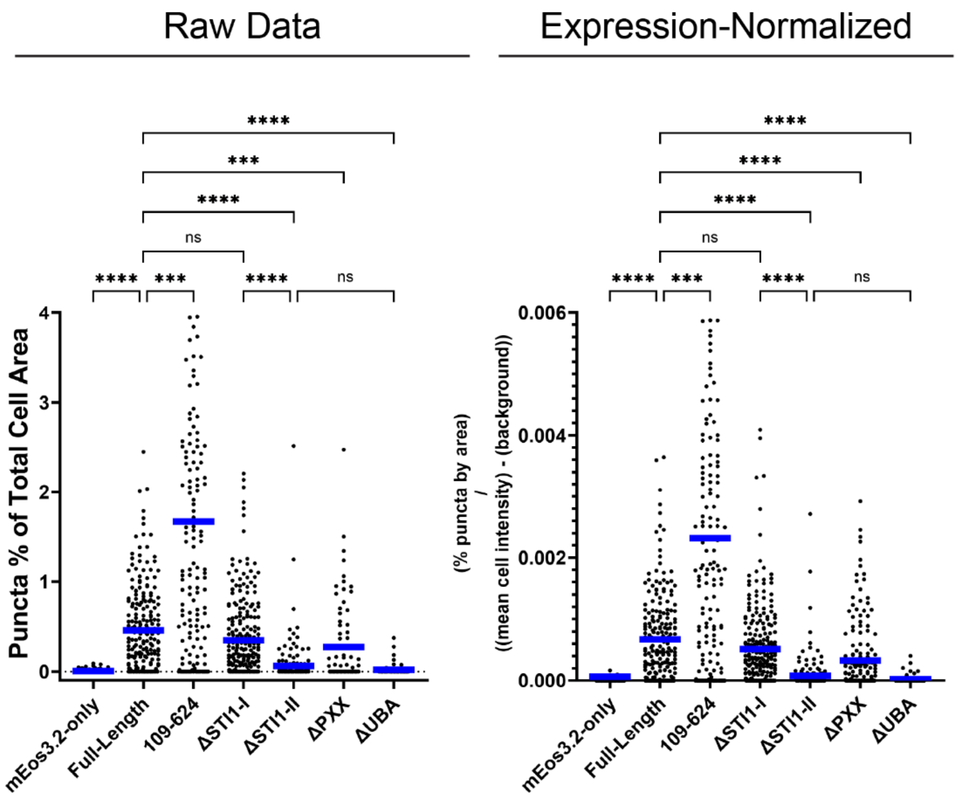


**Figure S3. Expression level does not account for differences in puncta formation between mEos3.2-UBQLN2 constructs.** Left: raw puncta data (puncta as a percentage of total cell area) from analyzed cells, reproduced from Figure 3. Right: the same puncta measurements normalized by expression level in each cell, calculated as (% puncta by area) / ((mean cell intensity) – (background)). Blue bars represent mean values. *p < 0.05, **p < 0.01, ***p < 0.001, ****p < 0.0001; ns, not significant; Kruskal–Wallis test with Dunn's multiple comparisons correction.

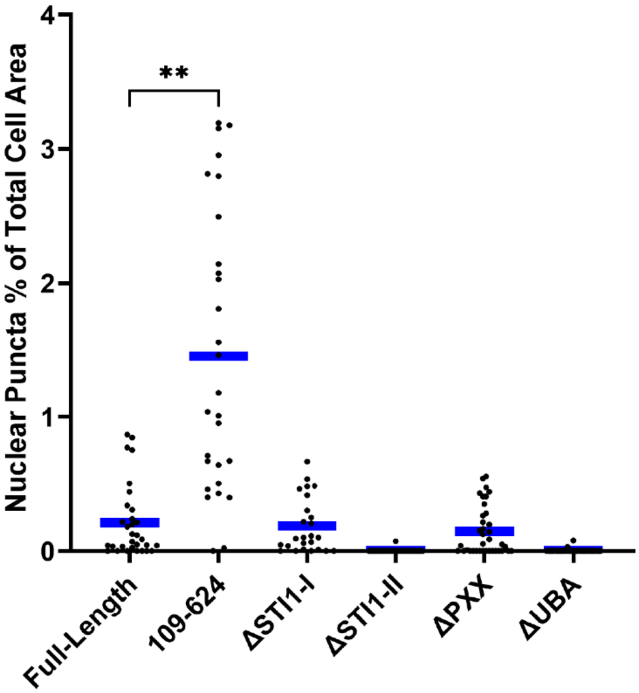


**Figure S4. Deletion of residues 1–108 leads to formation of nuclear UBQLN2 puncta.** Quantification of nuclear puncta as a percentage of total cell area for mEos3.2-UBQLN2 constructs: full-length, 109-624, ΔSTI1-I, ΔSTI1-II, ΔPXX, and ΔUBA. Data points represent individual cells from pooled experiments. Blue bars represent mean values. **p < 0.01; Kruskal–Wallis test with Dunn's multiple comparisons correction. n >= 48 cells per construct.

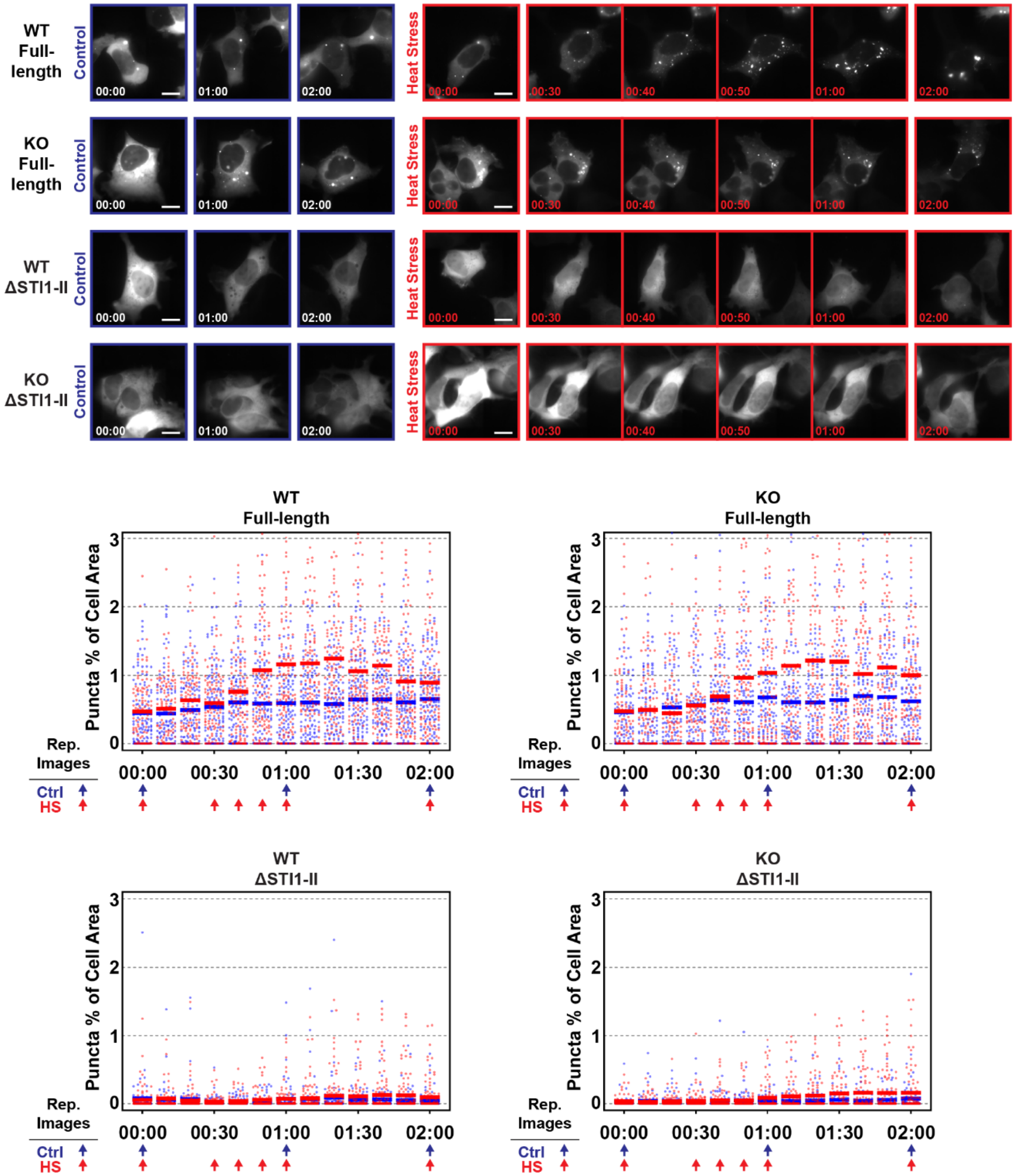
WT Full-length
KO Full-length
WT ΔSTI1-II
KO ΔSTI1-II
Control
Heat Stress
00:00
00:30
00:40
00:50
01:00
02:00
WT Full-length
KO Full-length
WT ΔSTI1-II
KO ΔSTI1-II
Puncta % of Cell Area
0
1
2
3
Rep. Images
Ctrl
HS
00:00
00:30
01:00
01:30
02:00

**Figure S5. mEos3.2-UBQLN2 puncta formation is not affected by knockout of endogenous UBQLN2.** Live-cell imaging of full-length mEos3.2-UBQLN2 and mEos3.2-UBQLN2 ΔSTI1-II in wild-type and *UBQLN2* knockout HEK293 cells under control (37°C) and heat stress (37°C to 42°C ramp) conditions. Top: puncta as a percentage of cell area over time for each condition, with individual cells plotted as data points and horizontal bars representing means. Control (blue) and heat stress (red) data are shown. Bottom: puncta percentage as a function of mean cell intensity for wild-type and knockout lines expressing each construct under control and heat stress conditions.

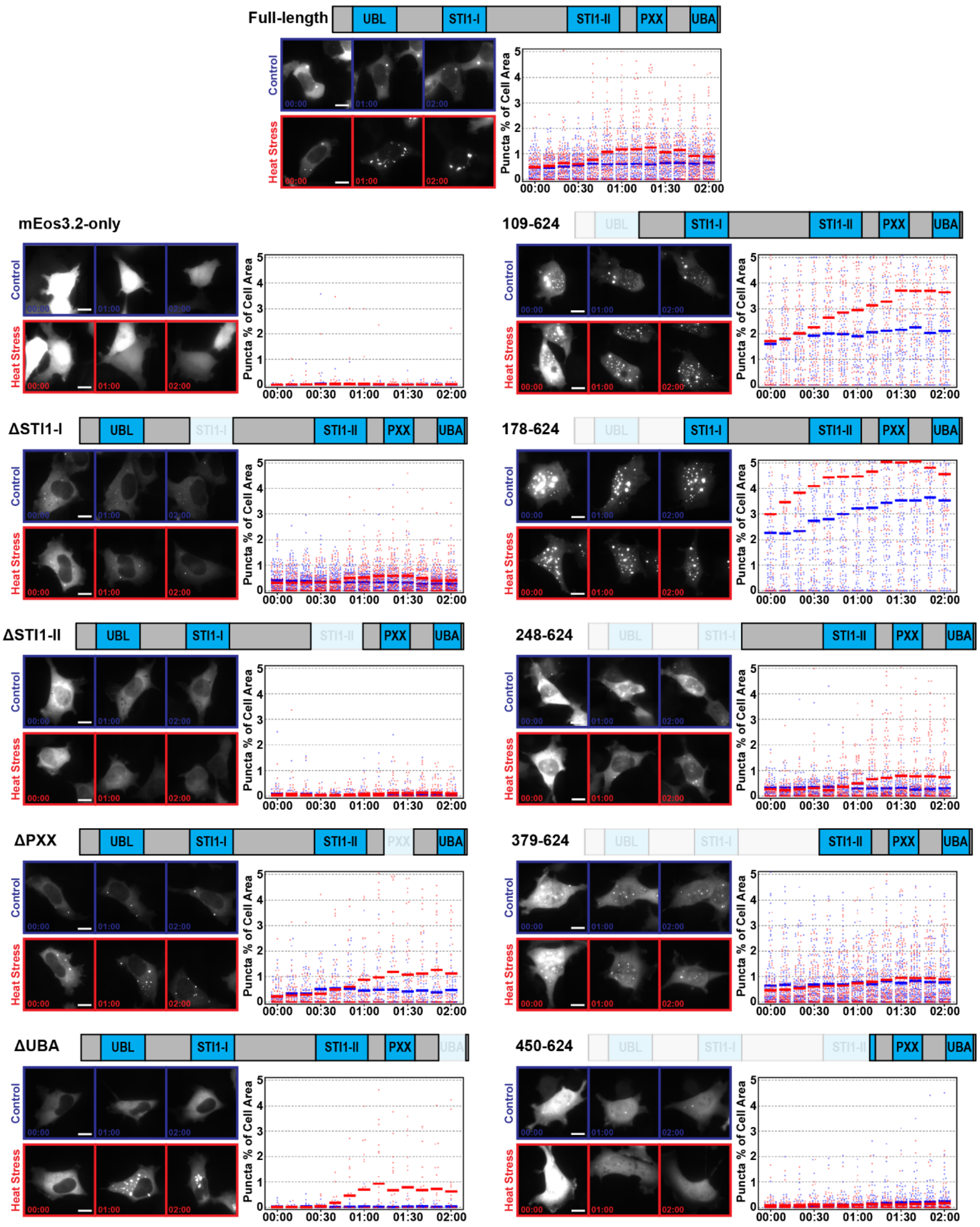

Full-length
UBL
STI1-I
STI1-II
PXX
UBA
mEos3.2-only
109-624
ΔSTI1-I
178-624
ΔSTI1-II
248-624
ΔPXX
379-624
ΔUBA
450-624
Control
Heat Stress
00:00
01:00
02:00
Puncta % of Cell Area
00:00 00:30 01:00 01:30 02:00

**Figure S6. Puncta quantification from live-cell heat stress timelapse experiments with UBQLN2 deletion construct library.** Comprehensive live-cell imaging data for all mEos3.2-UBQLN2 constructs subjected to control (37°C) and heat stress (37°C to 42°C ramp) conditions over 2 hours. Domain schematics are shown above each construct panel. Left column: single-domain deletion constructs (full-length, mEos3.2-only, ΔSTI1-I, ΔSTI1-II, ΔPXX, ΔUBA). Right column: N-terminal truncation constructs (109-624, 178-624, 248-624, 379-624, 450-624). For each construct, representative images are shown at 0, 1, and 2 hours under control (blue border) and heat stress (red border) conditions.

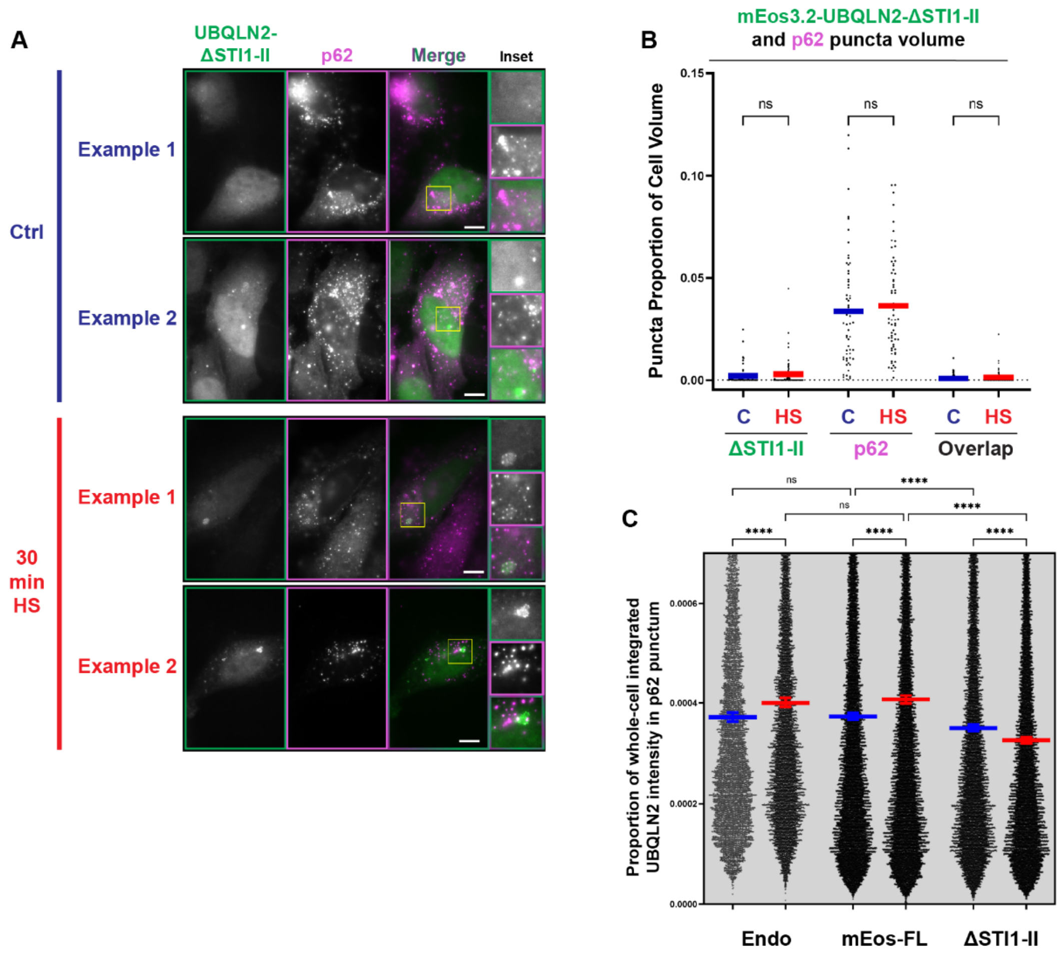


**Figure S7. Little or no enrichment of UBQLN2 ΔSTI1-II is observed in p62 puncta. (A, B)** Representative images and puncta volume quantifications from HeLa cells transfected with mEos3.2-UBQLN2 ΔSTI1-II, subjected to 37°C control or 30 min 42°C heat stress, then immunolabeled for endogenous p62 (magenta). Scale bars = 10 µm. Insets show raw signal. Graphs display the proportion of analyzed cell volume classified as puncta for UBQLN2 and p62 individually, as well as the overlapping proportion. C, control; HS, heat stress. **p < 0.01, *p < 0.05; ns, not significant; Mann-Whitney test. **(C)** Proportion of total integrated mEos3.2-UBQLN2 fluorescence intensity within spatially delimited individual p62 puncta for endogenous UBQLN2 and full-length and ΔSTI1-II UBQLN2 constructs under control and heat stress conditions. Each data point represents an individual p62 punctum. Horizontal bars represent the mean. ns, not significant; **p < 0.01, ****p < 0.0001, Kruskal–Wallis test with Dunn's multiple comparisons correction. C, control; HS, heat stress.

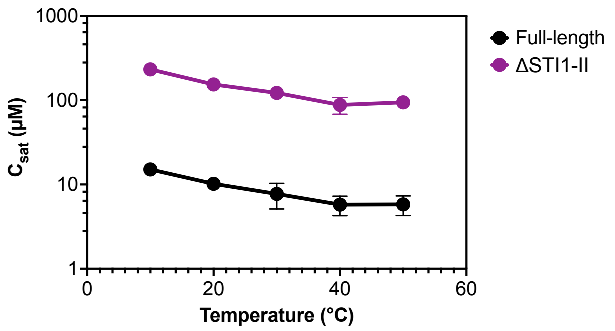


**Figure S8. ΔSTI1-II undergoes phase separation in the presence of crowding agent PEG8000, and not to similar extent as full-length UBQLN2.** Saturation concentration ($c_{sat}$) as a function of temperature for the indicated UBQLN2 constructs, determined by sedimentation assays with proteins expressed and purified from *E. coli*. Data points represent mean ± SEM across replicates.

### Supplementary Movies

**Movie S1.** An annotated compilation of live-cell experiments for mEOS3.2-UBQLN2 constructs. Sample movies for transfected HEK293 cells under control and heat stress conditions are provided.
